# Recurrent astrocyte transcriptional states with spatial and functional specialization are induced by brain metastases independently of their primary tumor origin

**DOI:** 10.64898/2026.11.16.766881

**Authors:** Carolina Hernández-Oliver, Lluís Cordón-Barris, Ana Sartorius-Yáñez, María González, Davide Maspero, Marc Elosua-Bayés, Neibla Priego, Leticia Cuarental, Laura Álvaro-Espinosa, Oliva Sánchez, Violeta Fernández, Guillermo Palou-Márquez, Carlos Parra-Pérez, Antonia Stammberger, Aránzazu Rosado, Ana Dopazo, María Ángeles Moro, Marc Schmitz, Holger Heyn, RENACER, Fátima Al-Shahrour, Manuel Valiente

**Author notes:** A list of authors appears at the end of the paper.

## Abstract

Reactive astrocytes have emerged as important components of the brain metastasis niche, yet a state-resolved framework for metastasis-associated astrocyte biology is still lacking. Here, we integrate single-cell and spatial transcriptomics across syngeneic breast, lung and melanoma models of brain metastases to define a state-resolved astrocyte landscape. We identify recurrent tumor-associated astrocyte states shared across tumor origins, map their spatial distribution, and find that a subset, which we term AST-BrM, becomes selectively reprogrammed at the tumor-microenvironment interface, supporting niche-coupled organization rather than a diffuse brain-wide reactive response. AST-BrM defines distinct metastasis-associated astrocyte states at the peritumoral interface, distinguishes them from homeostatic-like astrocytes beyond generic inflammatory activation, and resolves into two therapeutically actionable programs: an AP-1-high axis and a STAT3^+^/HOPX^+^-associated axis. Genetic and pharmacological perturbations establish functional relevance, and an AP-1/c-FOS-driven signature is associated with clinical outcome. Together, these findings define conserved, spatially organized and actionable astrocyte programs that drive brain metastasis progression.

## Introduction

Brain metastases are a frequent and clinically devastating complication of systemic cancer, with limited durable responses to current therapies and a strong dependence on the organ-specific microenvironment for successful colonization and outgrowth^1–3^. Beyond tumor-cell-intrinsic evolution, metastatic fitness in the brain is shaped by stromal circuits that regulate vascular co-option, immune remodeling, survival signaling and therapeutic response^4–6^, with astrocytes emerging as central regulators of this niche^7–12^. Specifically, mechanistic studies have shown that astrocytes can support brain metastasis through tumor-promoting signaling, inflammatory crosstalk, immune remodeling and resistance-associated programs^7,8,10–12^. However, these studies have largely defined individual circuits, often in the context of a single tumor origin, rather than a broader organizational framework for metastasis-associated astrocyte biology. In addition, brain metastasis progression is itself temporally structured: early colonization imposes strong microenvironmental constraints, particularly in the perivascular niche, whereas later stages involve progressive niche remodeling that supports metastatic outgrowth. This suggests that astrocyte functions are likely to evolve across disease stage rather than fixed throughout progression^4^.

This limitation is increasingly relevant because astrocyte identity and reactivity are now understood to be heterogeneous and context-dependent, rather than represented by a single stereotyped reactive state^13,14^.

Even in the healthy Central Nervous System (CNS), astrocytes are not uniform but are regionally specialized cell populations shaped by developmental patterning and local tissue context, giving rise to distinct transcriptional identities across anatomical regions^15–18^. Emerging evidence further suggests that these region-conserved astrocyte subgroups may represent distinct functional units, with differential capacities to support local circuit functions such as synaptogenesis or to engage disease-associated programs, including tumor pathology^19,20^. This intrinsic heterogeneity becomes further remodeled in disease, where multiple context-dependent astrocyte states have been described across inflammatory, ischemic and neurodegenerative conditions, arguing against a single generic reactive phenotype^14,21–23^. Together, these studies support the view that astrocyte responses are organized by local tissue context, disease stage and underlying regulatory program^24^.

In brain metastasis, however, it remains unclear whether astrocyte programs converge across tumor origins, how they are spatially organized within lesions, and whether distinct regulatory logics and vulnerabilities can be resolved in a manner that generalizes across models and cohorts. Recent work defining distinct tumor architectures and initiation microenvironments further supports the need for spatially informed, stage-resolved maps of astrocyte states in metastatic disease^25^.

Here, we address these gaps by constructing a state-resolved atlas of brain metastasis-associated astrocytes across syngeneic breast, lung and melanoma models, integrating single-cell profiling with spatial transcriptomics to map astrocyte programs onto tissue context. We derive a portable metastasis-associated astrocyte programs score (AST-BrM) that distinguishes tumor-associated from homeostatic-like astrocyte programs beyond generic inflammatory activation and enables cross-platform projection. We then resolve stage-linked regulatory logic, identifying a late-stage AP-1-centred program that is partially separable from canonical STAT3-associated reactivity, and test therapeutic tractability through genetic and pharmacological perturbation. Finally, we examine human relevance by anchoring conserved programs in an external NSCLC brain metastasis dataset^26^, validating tumor-adjacent AP-1 markers at the protein level, and assessing the clinical association of an AP-1/cFOS-linked signature in patient cohorts. Together, this work provides a unified framework for astrocyte heterogeneity in brain metastasis and identifies actionable niche programs that shape metastatic progression.

## Results

### Brain metastasis induces recurrent astrocyte states across primary tumor origins

Given the functional diversity and therapeutic relevance of astrocyte programs in brain metastasis^7,8,10,10–12^, we asked whether distinct tumor origins converge on shared astrocyte states. We profiled ACSA2^+^-enriched astrocytes from syngeneic breast (E0771-BrM^27^) and lung models (482N1^4^) (Fig. 1A), together with the melanoma brain metastasis model (B16/F10-BrM^9^) from Priego *et al.,* 2025. 7194 high-confidence astrocytes were retained after stringent annotation (Fig. 1A,B; Suppl. Fig S1A,B^28^). Unbiased clustering and state-specific signature enrichment (see Methods) identified 12 transcriptional states across models (AST1-AST12, Fig. 1C). Compared with healthy tissue, metastatic brains showed consistent remodeling, with AST5 emerging selectively in BrM, AST7, AST10, AST11 and AST12 enriched across brain metastasis models and AST6 preferentially represented in healthy brain (Fig. 1C; Suppl. Fig S1C, D). Hence, suggesting that brain metastasis induces a defined set of late-stage reactive programs rather than a single uniform reactive phenotype.

**Figure 1.**
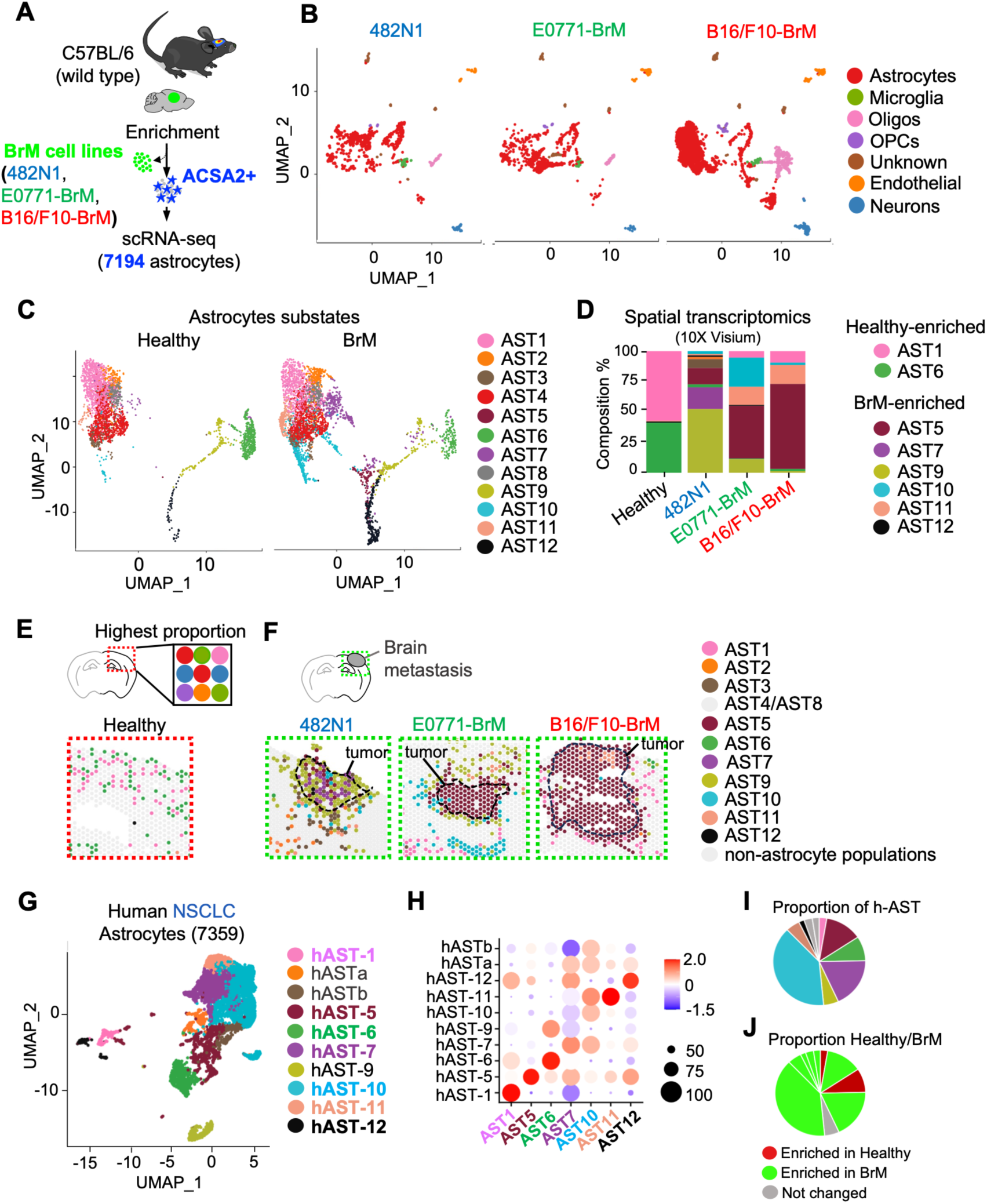
Brain metastasis induces recurrent astrocyte states across primary tumor origins. **A.** Schematic overview of the experimental design. Syngeneic brain metastasis models from lung (482N1) and breast (E0771-BrM) were established in C57BL/6 mice by intracranial injection. For the 482N1 and E0771-BrM models, three brains per condition were enzymatically digested and pooled. ACSA2 labeling was used to enrich in glial cells prior to scRNA-seq. The melanoma model (B16/F10-BrM) and control samples were obtained from Priego et al., 2025. A total of 7194 astrocytes were retained for downstream analyses. **B.** Uniform manifold approximation and projection (UMAP) embeddings of all captured cells for each model, shown separately and colored by manually annotated major cell class. Each model was analyzed and annotated independently. Melanoma model was reanalyzed in this study and large oligodendrocyte and microglia/macrophage populations were omitted from this visualization for clarity. Only populations found across all models are shown. **C.** Astrocyte-only UMAPs from healthy versus BrM conditions after signature-enrichment analysis (see Methods), resolving 12 astrocyte states (AST1-AST12). **D.** Barplot showing relative proportions of astrocyte-states across healthy tissue and BrM models inferred by deconvolution of spatial transcriptomics (10X Visium) data. States AST4 and AST8, whose abundance remained stable across conditions, were omitted for clarity (see Suppl. Fig S1F). Legend specifically divided into Healthy and BrM populations. **E,F.** Visium deconvolution maps showing the dominant (highest-proportion) astrocyte-state assignment per spot. Only zoomed-in regions of the sections are shown. Non-astrocyte populations are depicted in grey for better visualization. Also AST4/AST8 (see Suppl. Fig. 1H). Tumour regions are outlined in black. Red dashed boxes indicate healthy sample (**E**), whereas green dashed boxes indicate BrM models (**F**). **G.** UMAP of re-analysis of human NSCLC brain metastasis astrocytes (n=7359 cells) from Tagore *et al.,* 2025 identifying transcriptionally related human astrocyte states (hAST) and their correspondence to mouse astrocyte states. hASTa and hASTb were exclusively found in this cohort. **H.** Cross-species correspondence between human (rows) and mouse (columns) astrocyte states shown as a dot plot. Dot color indicates correlation/enrichment score, and dot size indicates the relative representation/overlap of each state. Warmer colors indicate higher correspondence, whereas cooler colors indicate lower correspondence. **I.** Pie chart showing the proportion of h-AST annotated in **G.** Legend as in **G. J**. Pie chart showing the proportion of Healthy and BrM-murine populations recapitulated in **H.**

Matched Visium spatial transcriptomics samples^29^ (Suppl. Fig. S1E) confirmed these shifts in tissue context. Projection of scRNA-seq-defined states with SPOTlight algorithm (Elosua-Bayés et al., 2021; Suppl. Fig. 1F) showed that healthy tissue was dominated by homeostatic states AST1 and AST6, whereas BrM samples displayed increased contribution of BrM-enriched states across tumor origins. Although AST4 and AST8 appeared relatively abundant in specific models (Suppl. Fig. S1G,H), their spatial distribution was broad and not preferentially associated with tumor regions (Suppl. Fig. S1G,H), in contrast to the BrM-enriched states, which preferentially occupy tumor-associated and peritumoral areas (Fig. 1D-F, Suppl. Fig S1H). Together, these data indicate that recurrent astrocyte programs are recruited or induced by tumor presence and become spatially organized within the metastatic niche, consistent with potential functional specialization, in line with mechanistic evidence that tumor-proximal astrocytes engage pro-metastatic circuits^5,6,9^.

Re-analysis of human NSCLC brain metastasis scRNA-seq^26^ recovered several murine-defined states, including AST1, AST5, AST6, AST11 and AST12 (Fig. 1G), while AST7 and AST10-like states appeared more heterogeneous in patients (Fig. 1H), suggesting that these programs may comprise additional substructure in patients and motivating deeper resolution in larger cohorts. Interestingly, the human astrocyte compartment was predominantly composed of BrM-associated states, whereas homeostatic-like states were comparatively underrepresented (Fig. 1I,J), consistent with the predominance of metastasis-associated reactive programs in astrocytes from established human brain metastases.

Together, these data indicate that brain metastasis induces recurrent astrocyte states across primary tumor origins with key components recapitulated across in human brain metastasis datasets.

### An AST-BrM signature defines a niche-coupled and functionally diverse astrocyte program in brain metastasis

To identify the populations most transcriptionally altered by brain metastasis we compared healthy and BrM-associated astrocytes. Differential expression followed by hierarchical clustering of the top differentially expressed genes separated astrocyte populations into two reciprocal transcriptional modules (Fig. 2A, Suppl.Table1,2), defining a healthy-enriched “AST” signature and a metastasis-associated “AST-BrM” signature (Fig. 2A, B). AST-BrM nominated AST5, AST7, AST10 and AST12 as the main BrM-associated states, as they contributed most strongly to the metastasis-enriched module (Fig. 2A) and were selectively expanded in BrM samples (Fig. 2B, Suppl. S1C-D). In contrast, AST healthy-enriched states were depleted or unchanged in BrM, consistent with a more homeostatic-like transcriptional profile (Fig. 2A-B, Suppl. S1C-D). Moreover, established astrocyte signatures confirmed enrichment of homeostatic programs in AST1, AST4 and AST6, versus A1-like inflammatory reactivity^14^ signature enrichment across the AST-BrM compartment (Suppl. Fig. S1I,J). However, AST9 displayed A1-like reactivity without AST-BrM enrichment, indicating that AST-BrM is not explained by generic inflammatory activation alone and supporting AST-BrM as a more metastasis-associated astrocyte program (Fig. 2A; Suppl. Fig. S1I,J).

**Figure 2.**
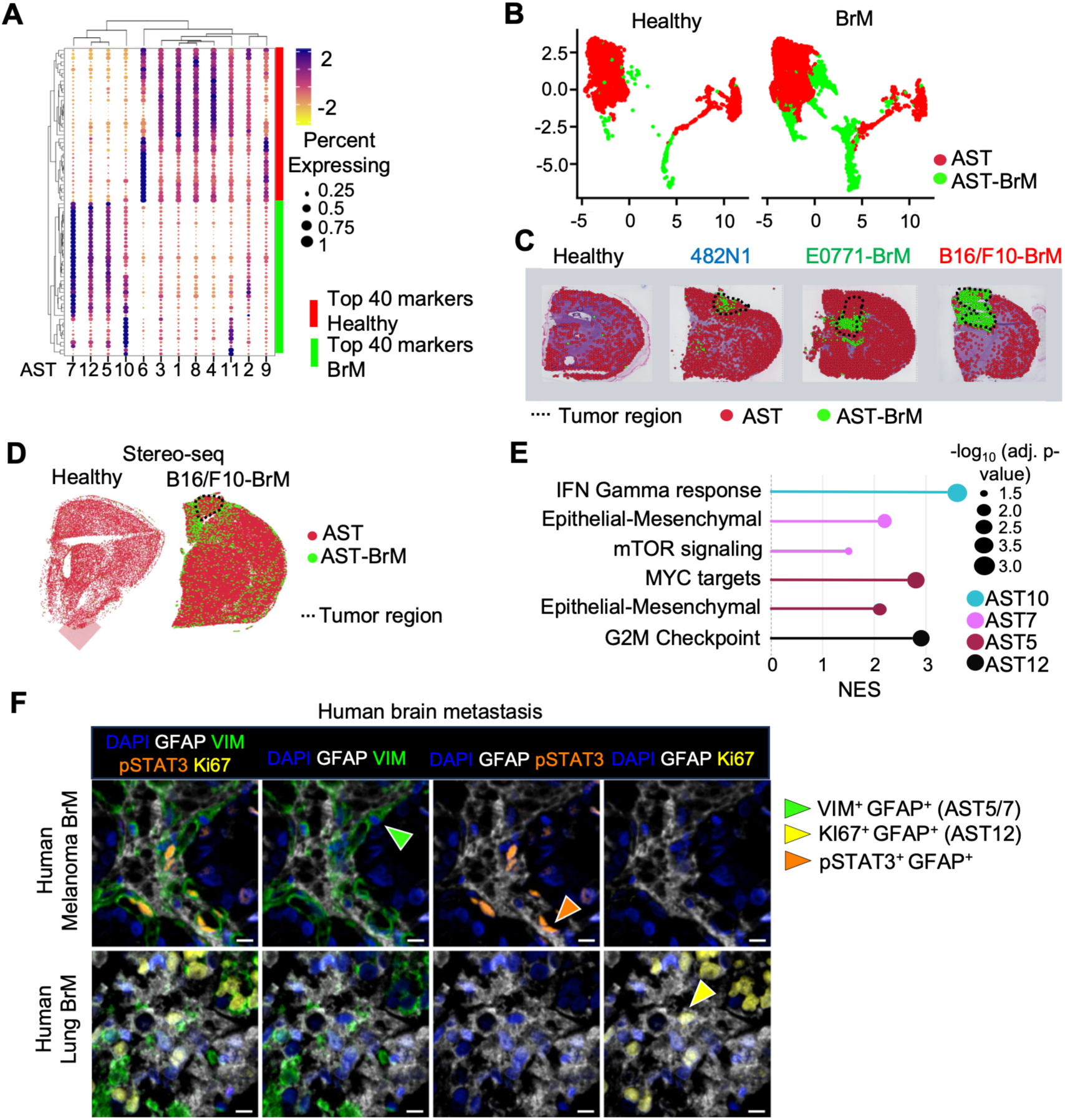
An AST-BrM signature defines a niche-coupled and functionally diverse astrocyte program in brain metastasis. **A.** Clustered dotplot of the top 40 differentially expressed genes between BrM and healthy astrocytes. Hierarchical clustering separates astrocytes states into a BrM-enriched module and a healthy-enriched module. Gene sets derived from these modules define the AST-BrM (BrM-enriched) and AST (healthy-enriched) signatures. Dot color indicates scaled average expression and dot size indicates percent of cells expressing each gene; side bar denotes astrocyte states (AST1–AST12). **B.** UMAP of astrocyte compartment for healthy and BrM conditions. AST populations (AST1, AST2, AST3, AST4, AST6, AST8, AST9, AST11) are labeled in red and AST-BrM populations (AST5, AST7, AST10, AST12) labeled in green. **C.** Spatial transcriptomics (10X Visium) deconvolution results highlighting AST populations in red (AST1, AST2, AST3, AST4, AST6, AST8, AST9, AST11) and AST-BrM populations in green (AST5, AST7, AST10, AST12) across samples (healthy, 482N1 lung-BrM, E0771-BrM breast, B16/F10-BrM melanoma). Tumor regions are outlined in black. **D.** Stereo-seq validation of the spatial organization of AST and AST-BrM signature scores in healthy brain and B16/F10-BrM samples. Gene-set enrichment analysis using the AST and AST-BrM signatures shows preferential enrichment of the AST-BrM program within tumor-associated regions. **E.** Gene set enrichment analysis across AST-BrM-driven states (AST5/AST7/AST10/AST12) reveals pathway-level separation, including immune-associated programs (e.g., IFN-γ response), proliferative signatures (G2/M checkpoint), and remodeling/metabolic axes (e.g., MYC targets, mTOR signalling, EMT-like programs). Dot size denotes −log10 adjusted p-value; x-axis shows normalized enrichment score (NES). **F.** Multiplex immunohistochemistry staining for GFAP, VIM, KI67, and pSTAT3 in human brain metastasis, showing distinct proliferative/reactive astrocyte phenotypes corresponding to AST5/7 (green) and AST12 (yellow) states. Arrowheads indicate representative cells for each population. *Scale bar, 10 μm. Sample from melanoma and lung primary human brain metastasis are used for these images (IDs B2347 for* melanoma BrM*; B22124 for* lung BrM*; Suppl.Table4)*.

Consistently, spatial projection of AST-BrM signatures at state resolution show specificity in tumor-associated regions, supporting niche-coupled engagement of distinct reactive programs rather than a uniform brain-wide response (Fig. 2C). Higher-resolution stereo-seq (Chen, A. et al., 2022) in the B16/F10-BrM model confirmed the peritumoral organization of AST-BrM astrocytes *in situ*, allowing a cross-platform validation (Fig. 2D).

We then performed gene set enrichment analysis to characterize the functional diversity within AST-BrM states. This revealed pathway-level separation, including immune-associated programs, proliferative signatures, and remodeling/metabolic axes such as IFN-γ response, G2/M checkpoint, MYC targets, mTOR signaling and EMT-like programs (Fig. 2E). Given previous evidence that a subset of RA in BrM is characterized by STAT3 activation, multiplex immunohistochemistry in human brain metastasis was used to examine how these newly defined AST-BrM phenotypes relate to the known GFAP^+^ pSTAT3^+^ compartment. This analysis confirmed the coexistence of distinct GFAP⁺ astrocyte phenotypes within the metastatic niche, including VIM⁺ remodeling-associated astrocytes corresponding to AST5/7 and Ki67⁺ proliferative astrocytes corresponding to AST12, both diverging from the established GFAP⁺ pSTAT3⁺ reactive astrocyte population^9^ (Fig. 2F). The distinct spatial and molecular profiles of these astrocyte subsets support the view that AST-BrM comprises multiple coexisting reactive programs engaged by different astrocyte populations in response to BrM.

Together, these data define AST-BrM as a spatially organized and functionally diverse metastasis-associated astrocyte program with potential relevance to brain metastasis progression.

### AST-BrM defines a late-stage AP-1-associated astrocyte response that diverges from canonical STAT3 reactivity

To nominate regulators of AST-BrM, we performed transcription factor motif enrichment analysis on the BrM-enriched AST-BrM gene set (RcisTarget^30–32^), NES > 3). This analysis revealed a strong over-representation of AP-1 family motifs (JUN/FOS branches and associated regulators) in AST-BrM astrocytes (Fig. 3A), consistent with AP-1-centred regulatory footprint in BrM-associated reactive astrocytes. To test whether this transcriptional signal translates *in situ*, we validated AP-1 activation at the protein level, detecting tumor-adjacent astrocytes positive for c-JUN and c-FOS in different BrM primary origins (Fig. 3B, C).

**Figure 3.**
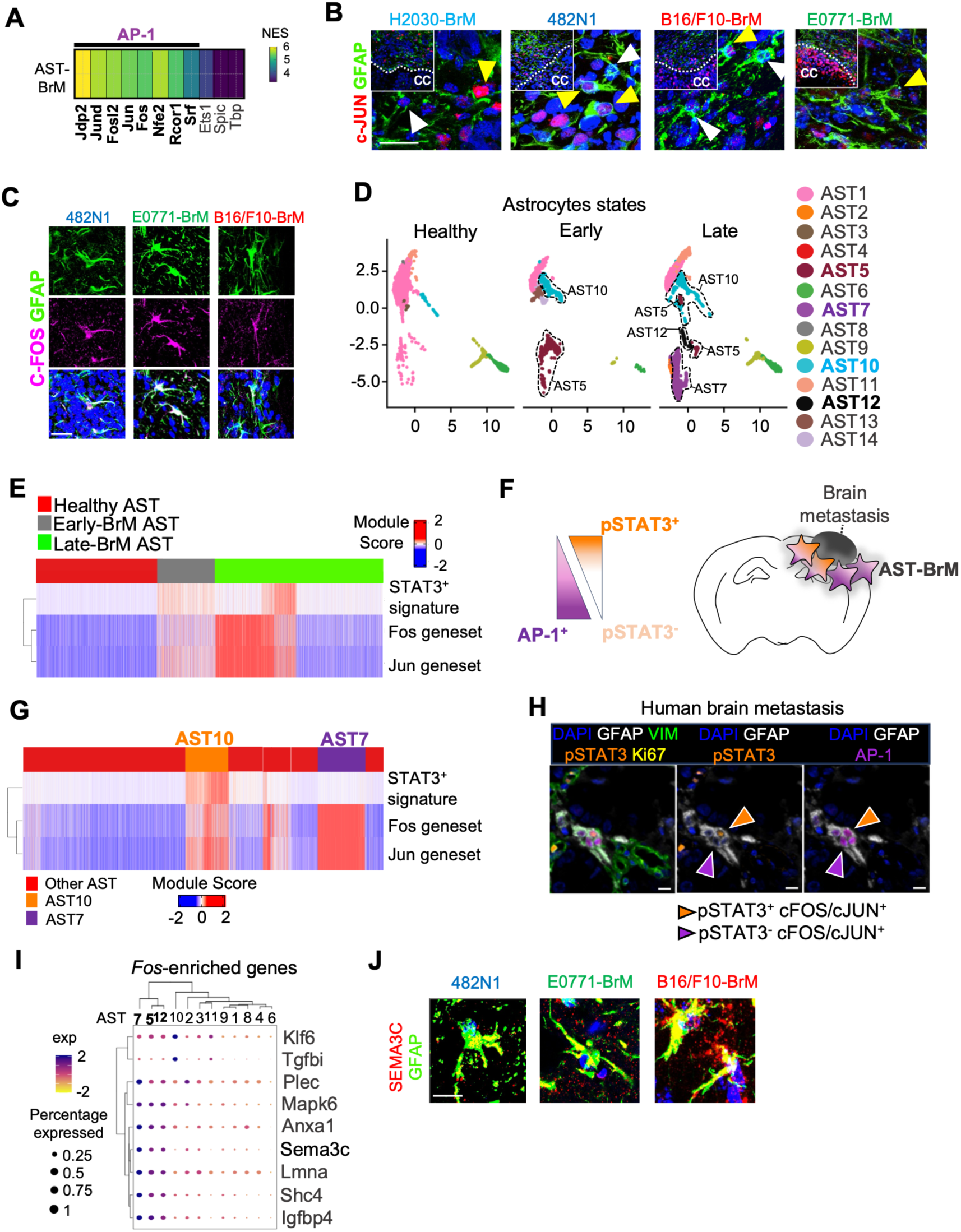
AST-BrM shows late-stage activation and bifurcates into AP-1- and STAT3-associated regulatory programs. **A.** Heatmap showing normalized enrichment scores (NES) of the top transcription factor regulons identified within the AST-BrM program by motif enrichment analysis with Scenic in the late-AST-BrM astrocytes. Strong enrichment of AP-1-related factors, including Jdp2, Jund, Fosl2, Jun and Fos is shown. **B.** Immunofluorescence staining for GFAP and c-JUN in brain metastases from the indicated models (H2030-BrM lung human-derived, 482N1 lung, B16/F10-BrM melanoma, E0771-BrM breast), showing c-JUN^+^ Ras at the peritumoral area. Heterogeneous engagement is shown across models (482N1 lung, B16/F10-BrM melanoma, E0771-BrM breast), meaning that not all RA are c-JUN^+^. Arrowheads indicate representative cells (doble positive in yellow; cJUN^-^astrocytes in white). CC, cancer cells. Scale bar, 75 μm. High magnification 20 μm. **C**. Immunofluorescence staining for GFAP and c-FOS in brain metastases from the indicated models, showing c-FOS-positive reactive astrocytes associated with metastatic lesions. Scale bar, 30 μm. **D.** UMAP embedding of astrocyte states across healthy, early and late BrM conditions, showing progressive emergence of late-stage-enriched astrocyte states. **E.** Heat map of module scores for STAT3, Fos and Jun gene signatures across healthy AST, early-BrM AST and late-BrM AST populations, showing preferential enrichment of Fos/Jun programs in late-stage astrocytes. **F**. Schematic representation of the two major regulatory axes resolved within AST-BrM, corresponding to an AP-1-high branch and a pSTAT3-associated branch. **G**. Heat map of module scores across astrocyte states highlighting bifurcation into AP-1-associated states (AST10/AST5) and STAT3-associated states (AST12/AST7), relative to other astrocyte states. **H**. Multiplex immunohistochemistry image of a human brain metastatic lesion showing GFAP, VIM, KI67, pSTAT3 and AP-1 (cFOS/cJUN) staining highlighting tumor-adjacent GFAP^+^ astrocytes with AP-1 signal in either a pSTAT3^+^ or pSTAT3^-^context, consistent with coexisting AP-1 and STAT3-associated RA states. Arrowheads indicate representative cells. Scale bar 10 μm. **I.** Dot plot of Fos-enriched genes across late-stage astrocyte states, highlighting selective enrichment of Sema3c within the AP-1-associated branch. Dot size indicates the proportion of expressing cells and color indicates scaled average expression. **J.** Representative immunofluorescence images showing SEMA3C expression in GFAP⁺ tumor-adjacent astrocytes across 482N1, E0771-BrM and B16/F10-BrM models. Scale bar 20 μm.

Given the evidence that astrocyte programs undergo stage-dependent transition during brain metastasis progression^4,9,33^, we next asked when AST-BrM and its AP-1 regulatory footprint become prominent. We profiled the glial compartment at 3 days post injection (dpi) in the 482N1 intracranial model and integrated it with the previously profiled late-stage 10 dp1 dataset, including technical replicates (Suppl. Fig. S2A-D; Fig. 3D). Because reactive astrocytes are expected to be less abundant at 3 dpi, we used GFAP-CreERT2;Ai14(tdTomato) mice and FACS isolation to enrich astrocytes at the early stage (Suppl. Fig. S2A-B). Consistent with previous work^9^, immunofluorescence confirmed that pSTAT3^+^ RA are rare at 3 dpi and enriched at 10 dpi (Suppl. Fig. S2E), confirming that this experimental design captures an established hallmark of metastatic astrocyte reactivity^9,34^.

At 3 dpi, early astrocytes were dominated by homeostatic-like programs, including synaptic-support features, while a subset already displayed immune-associated reactive signatures (Suppl. Fig. S2F, G; Suppl. Table3). In the integrated time-course atlas (Suppl. Fig. S2H, Fig. 3D), AST-BrM engagement was predominantly a late-stage feature, with enrichment concentrated in the dominant late BrM-associated states. Notably, when we repeated motif enrichment in early astrocytes, AP-1 was not a top regulatory signal; instead, ETS/SRF/NR3C1-type motifs ranked among the leading hits (Suppl. Fig. S2I), indicating a stage-dependent shift in regulatory logic. Consistently, per-cell module scoring showed that JUN/FOS target programs increase from healthy to early and peak at late stage, where they are enriched within the AST-BrM compartment (Fig. 3E).

To determine whether the AP-1-centred late-stage AST-BrM program overlaps with the established STAT3⁺ reactive astrocyte compartment, we scored a published STAT3⁺ reactive astrocyte signature^9^ across BrM astrocytes. STAT3 program scores were maximal in late-stage BrM astrocytes but showed only partial overlap with AP-1 module scores at single-cell resolution (Fig. 3E-G), indicating that the late-stage BrM astrocyte response comprises at least two separable regulatory programs: an AP-1-high program represented by AST7, and a STAT3-associated program represented by AST10 (Fig. 3F, G). We further validated these programs in situ by multiplex immunohistochemistry across human and murine BrM samples, confirming distinct tumour-adjacent GFAP⁺ astrocyte subpopulations corresponding to the VIM⁺ AST5/7 and KI67⁺ AST12 states identified previously (Fig. 2E), together with AP-1^-^and/or pSTAT3-positive astrocytes consistent with the regulatory bifurcation defined in Fig. 3F (Fig. 3H). cFOS validation showed positive coexpression (Fig. 3C). To identify downstream features associated with the AP-1 high branch of AST-BrM, we examined Fos-enriched genes across late-stage astrocyte states. Among these, Sema3c emerged as an AP-1 associated marker expressed mostly by AST7 (Fig. 3I), and its expression was further validated *in situ* across BrM models (Fig. 3J).

Together, these data position AST-BrM as a late-stage-enriched astrocyte response marked by increasing AP-1 activity and partial divergence (Fig. 3H) from canonical STAT3-associated reactivity. This regulatory separation suggests that late BrM-associated astrocytes comprise coexisting AP-1- and STAT3-associated states within the metastatic niche, providing a framework to functionally interrogate these axes.

### Functional targeting STAT3-independent AP-1^+^ and STAT3-dependent HOPX^+^ astrocyte axes impairs brain metastasis viability

To test the functional contribution of AP-1^+^ RAs to brain metastasis, we generated conditional mutant mice lacking cFOS from RAs (cKO-cFos: *GFAP-Cre/ERT2*;*cFos^loxP/loxP^*) in which syngeneic B16/F10-BrM brain metastatic (BrM) cells, labeled with luciferase, were intracardially inoculated (Fig. 4A). These models have gained brain tropism while retaining an aggressive extracranial metastatic potential^9^. Three days after intracardiac inoculation of B16/F10-BrM in cKO-cFos mice, we injected tamoxifen (Tmx), and 13 days later, we analyzed brain metastasis with bioluminescence imaging (BLI) (Fig. 4B). A significant reduction in brain metastasis burden was detected in cKO-cFos mice treated with Tmx (n = 42, P= 0.0348) (Fig. 4B, C) demonstrating that c-Fos component of AP-1 complex in brain metastasis-associated astrocytes is required for disease progression (Fig. 4C). Importantly, because this intracardiac inoculation captures dissemination and brain colonization from the systemic circulation, this *in vivo* genetic perturbation orthogonally validates our transcriptional and spatial findings derived from intracranial models, reinforcing AP-1 RAs as a niche-relevant astrocyte dependency in brain metastasis.

**Figure 4.**
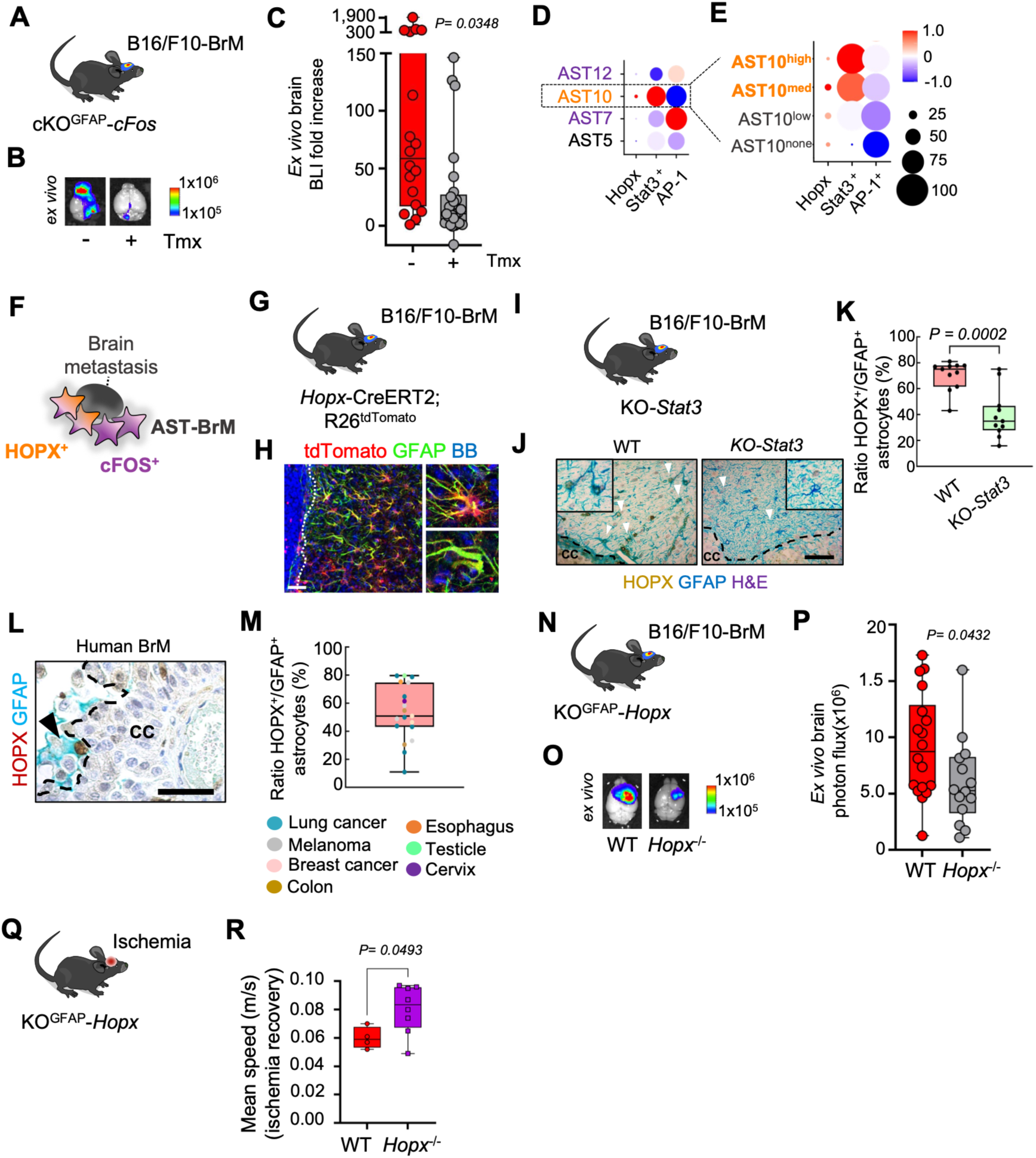
Functional targeting of AP-1 (cFOS) and HOPX astrocyte axes impairs brain metastasis and reveals context-dependent roles in ischemic recovery. **A.** Experimental design for astrocyte-specific deletion of cFOS. GFAP-Cre/ERT2; cFOS^loxP/loxP^ mice were injected intracardially with B16/F10-BrM cells and treated with tamoxifen (Tmx) to induce recombination. Ex vivo bioluminescence imaging (BLI) was used to assess metastatic burden. Tmx treatment was done intraperitoneal. **B.** Representative ex vivo bioluminescence images of cKO^GFAP^-cFOS mice in the absence (-) or presence (+) of tamoxifen-induced recombination 2 weeks after being inoculated with B16/F10-BrM cells intracardially. A total of 42 mice were included. **C.** Quantification of BLI. Values are shown in box-and-whisker plots where every dot represents a different animal and the line in the box corresponds to the median. Whiskers go from the minimum to the maximum value. Values were obtained from normalizing the ex vivo brain signal to the in vivo head signal 3 days after intracardiac injection when treatment was initiated (n = 21 mice per experimental condition, 2 independent experiments). P-value was calculated using the two-tailed t-test. **D.** Dot plot of the average expression of *Hopx*, *Stat3* and *cFos/cJun* AP-1 regulatory genes across AST-BrM populations (AST5, AST7, AST10, AST12). Dot size indicates the proportion of cells, and color (blue, red) indicates scaled enrichment score. Colors for letters in AST populations follow rules in Fig. 3F. **E.** Dot plot of the average expression of *Hopx, Stat3* and *cFos/cJun* AP-1 regulatory genes across AST10 subpopulations stratified by *Stat3* expression level: AST10^none^, AST^low^, AST10^med^, AST10^high^ denote AST10 cells with absent, low, intermediate and high *Stat3* expression, respectively. Dot size indicates the proportion of cells, and color (blue, red) indicates scaled enrichment score. **F.** Schema of the 2 different axes that govern AST-BrM programs. *Hopx*^+^ as *Stat3*-dependent and *cFos*^+^ as *Stat3*-independent. **G.** Experimental design for *Hopx*-expressing astrocytes in brain metastasis using Hopx-CreERT2; R26^tdTomato^ reporter mice. **H.** Representative immunofluorescence image showing tdTomato-labelled HOPX-lineage cells in a B16/F10-BrM lesion, co-stained for GFAP. **I.** Experimental scheme of the B16/F10-BrM model in cKO-STAT3 mice. **J**. Representative immunohistochemistry images showing HOPX and GFAP staining in WT and cKO-STAT3 brain metastases of B16/F10-BrM model. Arrowheads indicate HOPX^+^/GFAP^+^ astrocytes. Scale bar, 50 μm. **K.** Quantification of the proportion of HOPX^+^/GFAP^+^ astrocytes in WT and cKO-STAT3 lesions. P-value was calculated using the two-tailed t-test. **L.** Representative immunohistochemistry image of human brain metastasis showing HOPX^+^/GFAP^+^ astrocytes. Arrowheads indicate representative double-positive cells. Scale bar, 50 μm. **M.** Quantification of the percentage of HOPX^+^/GFAP^+^ astrocytes across human brain metastasis from multiple primary tumor origins. P-value was calculated using the two-tailed t-test. **N.** Experimental design for astrocyte-specific deletion of *Hopx*. GFAP-Cre/ERT2; *Hopx*^loxP/loxP^ mice were injected intracranially with B16/F10-BrM cells. Ex vivo bioluminescence imaging (BLI) was used to assess metastatic burden. **O.** Representative ex vivo bioluminescence images of KO^GFAP^-HOPX mice after being inoculated with B16/F10-BrM cells intracranially. n=18 WT; n=14 cKO mice were included. **P.** Quantification of BLI. Values are shown in box-and-whisker plots where every dot represents a different animal and the line in the box corresponds to the median. Whiskers go from the minimum to the maximum value. Values were obtained from the ex vivo brain signal at endpoint after intracranial injection (n = 33 mice, 1 independent experiment). P-value was calculated using the two-tailed t-test. **Q.** Experimental design for ischemia model. KO^GFAP^-HOPX model was used to induce ischemia. **R.** Quantification of mean speed during behavioural testing in WT and Hopx^−/−^ mice, showing altered functional recovery despite similar acute lesion size. P-value was calculated using the two-tailed t-test.

Based on our AST-BrM stratification, late-stage BrM-associated astrocytes segregated into two regulatory programs characterized by AP-1 activity and STAT3-associated reactivity, respectively. We therefore asked whether the STAT3-associated branch engages stemness-like features that could represent an additional functional layer beyond immunomodulation.

Prior work showed that STAT3^+^ RAs acquire stemness-like features in brain metastasis, including Nestin enrichment and enhanced astrosphere self-renewal, but the functional relevance of this property was not tested^9^. Because *Hopx* marks quiescent/stem-like neural stem cell populations and regulates germinal activity^35^ and was highly expressed in a subpopulation within the STAT3^+^ axis of our AST-BrM compartment (Fig. 4D-F), we tested whether astrocyte-specific *Hopx* perturbation impacts metastatic fitness. We first validated the HOPX enrichment in experimental brain metastasis (Fig. 4G,H) and found that its expression was reduced upon *Stat3* deletion, supporting HOPX as a downstream feature of the STAT3-associated astrocyte programme (Fig. 4I-K). We further confirmed HOPX^+^/GFAP^+^ astrocytes in human brain metastasis across multiple tumor origins (Fig. 4L,M).

We next generated conditional mutant mice lacking *Hopx*, specifically from RAs (cKO-Hopx: *GFAP-Cre/ERT2*;*Hopx^loxP/loxP^*). We performed an intracranial inoculation of B16/F10-BrM cell line in KO-Hopx mice (Fig 4N), and 10 days later, we analyzed brain metastasis with BLI. A significant reduction in brain metastastasis burden was detected in KO-Hopx mice (n= 33, P=0.0432) (Fig. 4O,P).

Given that *Hopx* has been implicated in broader CNS injury-associated glial responses^36^, we asked whether the *Hopx*-associated astrocyte program might also influence functional outcome after cerebral ischemia (Fig. 4Q). Specifically, to next assess whether astrocyte-specific *Hopx* deletion affects long-term functional outcome after cerebral ischemia, we evaluated spontaneous locomotor activity in the open field test one month after middle cerebral artery occlusion (MCAO, Suppl. Fig. 3A). Acute MRI assessment performed 48h after MCAO (Suppl. Fig. 3B) did not reveal genotype-dependent differences in infarct volume or edema between conditions (Suppl. Fig. 3B-D), indicating that *Hopx* deletion does not markedly alter the early extent of ischemic injury. In contrast, behavioral analysis at a later post-ischemic stage (Suppl. Fig. 3A) revealed improved locomotor performance in *Hopx*-deficient MCAO mice (Supp. Fig. 3E,F). In the open field test, *Hopx*-deficient mice displayed higher mean speed (Fig. 4R) suggesting that loss of *Hopx* promote motor recovery after ischemic injury. These results indicate that HOPX^+^ RA are a disease-promoting glial cell subpopulation beyond brain metastasis as it limits motor recovery during the chronic phase after ischemic injury.

Together, these findings define AST-BrM as a functionally relevant astrocyte program that supports brain metastasis through distinct AP-1/cFOS^+^ and HOPX-associated mechanisms, while implicating the HOPX axis in post-ischemic recovery.

### Pharmacological targeting AP-1^+^ astrocyte axis impairs brain metastasis viability

To assess the clinical relevance of the AP-1^+^ astrocyte program identified in our preclinical models, we first examined the RENACER^37^ bulk RNA-seq cohort with annotated clinical outcomes (Fig. 5A,B). We computed ssGSEA scores for a cFOS-enriched gene set, defined as the subset of AST-BrM genes annotated as putative cFOS/AP-1 regulon targets in SCENIC (Fig. 3K). Because bulk profiles integrate variable tumour purity and stromal composition, we stratified samples according to tumour cell content (Fig. 5B). Notably, high expression of the Fos-enriched signature was associated with worse overall survival selectively in high TME samples (Fig. 5D; n = 112; P=0.00041, HR=3.5, FDR<0.001), whereas no significant association was observed in low TME samples (Fig. 5C; n = 112; P=0.12, HR=1.5, FDR=0.2). Consistently, sensitivity analysis showed that the prognostic effect of the *cFos* program was strongest in low-purity samples (Fig. 5E), arguing that, although *cFos* is detectable in both tumour cells and reactive astrocytes (Fig. 5F-H), its prognostic value is preferentially driven by the tumor microenvironment rather than by a purely tumor-intrinsic transcriptional program. We next sought tissue-level evidence for AP-1 activation in human disease. We analyzed a retrospective set of 9 FFPE BrM samples previously curated in Priego *et al.,* 2025, selected from extended neurosurgical resections to ensure substantial peritumoral microenvironment was represented. Immunohistochemistry confirmed cFOS signal within GFAP⁺ astrocytes, consistent with AP-1 activation in tumor-adjacent astrocytes in human disease (Fig. 5F-H).

**Figure 5.**
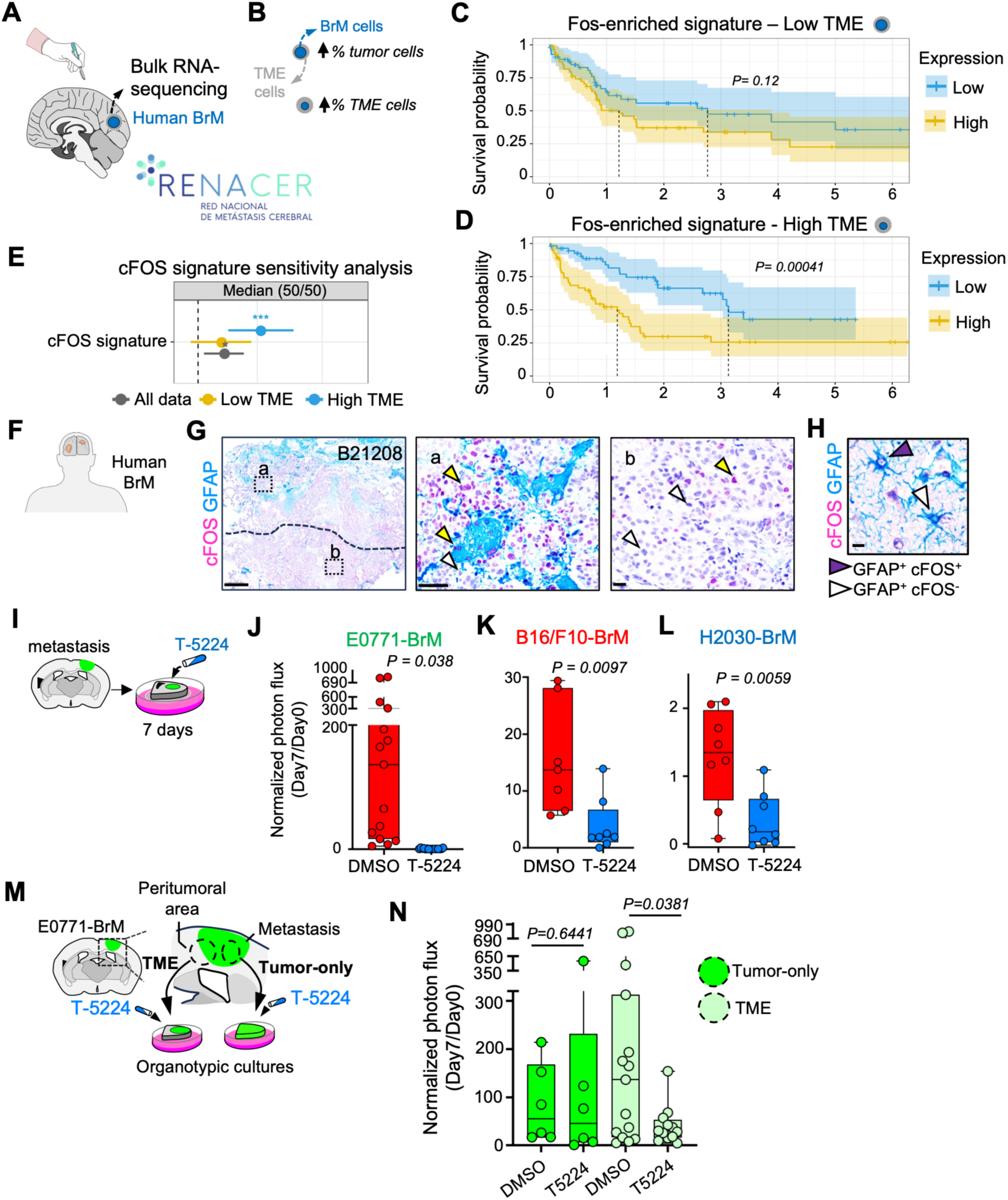
AP-1 astrocyte activation is clinically relevant and pharmacologically targetable in ex vivo brain metastasis. **A.** Schematic of the RENACER human brain metastasis cohort used for bulk RNA-seq analysis. **B**. Samples were stratified according to tumor purity to distinguish tumor-rich and TME-rich microenvironments. **C.D.** Kaplan-Meier analysis of overall survival according to expression of the Fos-enriched signature by ssGSEA score in tumour-rich **(C)** and tme-rich **(D)** samples. **E**. Sensitivity analysis of the prognostic association of the *cFos* signature across the full cohort and after purity stratification, showing stronger prognostic value in stroma-rich samples. **F.** Schematic of the analysis of human brain metastasis samples. **G.** Representative immunohistochemistry image showing cFOS and GFAP staining in a human brain metastasis, with higher-magnification views of the boxed regions (a) stroma-rich region (b) tumor rich region. **H.** Representative example of GFAP⁺ astrocytes with or without cFOS signal in the tumour-adjacent region. **I.** Experimental scheme of organotypic brain slice cultures treated with the AP-1 inhibitor T-5224 containing established intracranial metastatic lesions. **J-L.** Cross-model evaluation of T-5224 in organotypic cultures containing E0771-BrM (breast), B16/F10-BrM (melanoma) and H2030-BrM (lung, human-derived) lesions, showing reduced ex vivo tumour outgrowth upon AP-1 inhibition across models. Tumour burden quantified longitudinally by bioluminescence imaging (BLI) of luciferase-expressing metastatic cells. P-value was calculated using the two-tailed t-test. **M.** Compartment-controlled organotypic cultures generated by microdissection of tumour core only versus tumour plus peritumoral TME. **N**. Evaluation of T-5224 in Compartment-controlled organotypic cultures depicted in **J**. T-5224 produced a stronger reduction in tumour burden when the peritumoral microenvironment was retained, supporting a microenvironment-amplified component of AP-1–dependent metastatic fitness. P-value was calculated using the two-tailed t-test.

To next asses the pharmacological potential of targeting AP-1, we used T-5224, a small-molecule inhibitor that selectively blocks cFOS/cJUN DNA binding and suppresses AP-1-dependent transcription^38–40^. We first evaluated tumor-intrinsic sensitivity *in vitro* (Suppl. Fig. 4A) and then focused on organotypic brain slice cultures (Fig. 5I), which preserve brain cytoarchitecture and tumor-microenvironment interactions while enabling controlled pharmacological perturbation^41^. In organotypic cultures containing E0771-BrM lesions, T-5224 treatment reduced tumor outgrowth measured by bioluminescence (Fig. 5J), also across other models (Fig. 5 K, L) despite minimal effects on tumor cell number at the same concentration *in vitro* (Suppl. Fig. 4A-B). This separation between *in vitro* and *ex vivo* responses suggest that AP-1 inhibition impacts a microenvironment-enabled component of metastatic growth.

We therefore tested whether the tumor microenvironment (TME) modulates sensitivity to AP-1 inhibition by generating compartment-controlled organotypic cultures containing either tumor core alone or tumor plus peritumoral tissue (Fig. 5M). T-5224 produced a stronger reduction in tumor burden when the peritumoral TME was retained, whereas tumor-only cultures showed a weaker response at the same dose (Fig. 5M, N). Together, these data indicate that AP-1 blockade limits metastatic fitness through mechanisms that are amplified by tumor-brain niche interactions, consistent with our single-cell and spatial evidence that AP-1 activation is prominent in tumor-adjacent astrocytes. These findings, together with previous ones, nominate the AP-1 astrocyte axis as a pharmacologically actionable niche dependency in brain metastasis.

## Discussion

Our study challenges the long-standing paradigm that brain metastasis-associated astrocytes behave as a uniform population. By resolving astrocytes by tumor origin and disease stage, we show that their functional impact on the metastatic niche depends on cellular heterogeneity and that incorporating these axes is essential for defining therapeutically actionable astrocyte states. Importantly, we find that a recurrent set of astrocyte programs (named as AST-BrM) is induced across lung, melanoma and breast brain metastasis models, supporting the idea that metastatic cells, despite distinct origins, can converge on shared astrocytic responses. This observation aligns with and extends prior work implicating tumor-proximal astrocytes in distinct pro-metastatic circuits^5–12^ and reinforces the concept that brain metastasis engages multiple reactive programs rather than a single “reactive astrocyte” phenotype^42^. At the same time, the upstream drivers of this reprogramming remain incompletely defined. Whether convergent astrocyte states arise from shared tumor-derived cues, common tissue-injury signals, or coordinated multicellular interactions in the niche, will require systematic dissection. Together, our results support a model in which brain metastasis recruits a collection of distinct astrocyte subprograms that coexist within the tumor microenvironment. Consistent with this, AST-BrM engagement becomes most prominent during late-stage outgrowth, consistent with progressive recruitment or induction of tumor-associated reactive programs as lesions establish. Importantly, the regulatory logic also shifts across stages: AP-1 emerges as a dominant footprint in late-stage AST-BrM astrocytes, whereas early-stage astrocytes show different motif enrichments, supporting the idea that astrocyte programs are dynamically rewired during progression.

At the same time, our data support strong tissue and niche specificity. AST-BrM programs that associate with metastatic outgrowth were spatially confined to tumor-adjacent and peritumoral territories, consistent with regulation by local microenvironmental cues. This pattern suggests that short-range signaling and/or contact-dependent communication at the tumor-microenvironment interface may be important for inducing and maintaining these astrocyte states. This spatial organization is consistent with mechanistic studies demonstrating that tumor-proximal astrocytes engage pro-metastatic circuits, including exosome-mediated tumor reprogramming and gap-junctional communication^5,6^, as well as STAT3-driven immunomodulatory reactivity^10^. Together, our data extend these seminal mechanistic observations by positioning them within a unified state framework that generalizes across tumor origins.

AST-BrM is robustly transferable across platforms and cohorts, projecting into spatial datasets and showing that metastasis-associated astrocyte programs are anatomically compartmentalized within tumour-associated regions. Notably, AST-BrM is not explained by generic inflammation alone, as inflammatory scores can be elevated in states lacking AST-BrM enrichment. Human relevance is supported by recovery of multiple murine-defined programs in an external NSCLC-BrM dataset and by protein-level validation of AP-1 markers and HOPX in tumour-adjacent astrocytes. Together, these findings support conserved niche-associated astrocyte programs, while also indicating additional patient-specific heterogeneity that will require larger multi-origin cohorts to resolve fully.

A major strength of this work is the definition and functional testing of two partially separable astrocyte axes within the BrM niche. First, genetic perturbation of the AP-1 and STAT3/HOPX branches demonstrate causal contributions to metastatic fitness: astrocyte-specific deletion of cFOS reduced brain metastatic burden, and perturbation of the STAT3/HOPX-associated program similarly impaired metastatic outgrowth, supporting that these programs are not merely correlative state markers. Moreover, the context-dependent impact of Hopx deficiency on ischemic recovery underscores a broader principle: pro-metastatic astrocyte programs are embedded in general astrocyte biology of tissue remodeling and recovery, highlighting the need to define therapeutic windows and downstream effectors that minimize disruption of essential homeostatic functions. Complementing these genetic data, pharmacological AP-1 inhibition with T-5224 reduced metastatic outgrowth in organotypic brain slice cultures at concentrations with limited tumor-intrinsic effects *in vitro*, and compartment-controlled experiments showed that retention of the peritumoral microenvironment amplifies sensitivity to AP-1 blockade, consistent with a niche-enabled component of AP-1-dependent metastatic fitness^38,41^. Together, these findings support the existence of actionable, state-linked astrocyte dependencies organized along AP-1 and STAT3/HOPX axes and help explain why prior studies have identified distinct astrocyte-dependent mechanisms across contexts and tumor types^5–9^.

Several limitations of this study should be considered. Although we recover key features of AST-BrM in human datasets, the extent of inter-patient and cross-origin heterogeneity in human brain metastasis remains incompletely resolved. In addition, while our perturbation experiments establish functional relevance for the AP-1/cFOS and STAT3/HOPX axes, the downstream effectors through which these programs influence metastatic fitness remain to be defined. Future studies in larger human cohorts and with higher-resolution spatial approaches will therefore be important to refine the architecture and therapeutic vulnerabilities of metastasis-associated astrocyte states.

In sum, we propose that brain metastasis consistently remodels astrocytes into a recurrent, spatially organized state space in which late-stage tumor-adjacent programs are organized by at least two partially separable regulatory axes: AP-1 and STAT3/HOPX. This framework reconciles mechanistic diversity in the literature^5–9^, enables cross-platform detection through AST-BrM program scoring^43^, and identifies niche dependencies that can be targeted^38,41^.

## Methods

### Animal studies

All animal experiments were performed in accordance with protocols approved by the CNIO, Instituto de Salud Carlos III and Comunidad de Madrid Institutional Animal Care and Use Committee (PROEX299/15 and PROEX135/19). The cKO^GFAP^-*cFos* mice were generated by breeding GFAP-CRE/ERT2 mice [B6.Cg Tg(GFAP-cre/ERT2)505Fmv/J, The Jackson Laboratory, cat.# 012849] with cFOS*^l^*^oxP/loxP^, kindly shared by Dr. Erwin Wagner Enriquez. cKO^GFAP^-*Hopx* mice were generated by breeding GFAP-Cre/ERT2 mice described above with Hopx^loxP/loxP^ from Lin et al., 2020. For early-stage astrocyte profiling, GFAP-Cre/ERT2;Ai14(tdTomato) reporter mice were used to enable isolation of tdTomato-positive astrocytes.

Brain colonization assays were performed in C57BL/6 mice 6–10 weeks as previously described (Priego et al., 2018), by injecting 100 μL of PBS into the left ventricle containing 100,000 cancer cells or 1μL of DMEM plain medium intracranially (right frontal cortex, approximately 1.5 mm lateral and 1 mm caudal from the bregma, and to a depth of 2 mm) containing 40,000 cancer cells by using a gas-tight Hamilton syringe and a stereotactic apparatus.

Metastatic colonization was analyzed *in vivo* weekly and *ex vivo* at the endpoint by BLI. Anesthetized mice with isoflurane were injected with D-luciferin (150 mg/kg) retro-orbitally and imaged in an IVIS machine. Average radiance (photons/sec/cm^2^/steradian) from each region of interest (ROI) was measured using Living Image software, version 4.5. ROIs were created in the 2D bioluminescent image in the head or body of each mouse for *in vivo* analysis or around the corresponding organ for ex vivo analysis. For intracranial injections, *ex vivo* values at the end point were taken. For intracardiac injections, *ex vivo* values at the end point were normalized to the BLI values of the head *in vivo* 3 days after injection of the cancer cells.

Cre expression was induced by tamoxifen (Tmx) 3 days after intracardiac or intracranial inoculation of cancer cells, when it is considered that the first brain metastatic cells have crossed the blood brain barrier. AP-1 inhibitor (T5648, Sigma-Aldrich) was prepared as a 20 mg/mL stock solution in corn oil. Mice were administered tamoxifen via intraperitoneal (i.p.) injection at a dose of 50 mg/kg body weight. The administration schedule consisted of 5 consecutive days of tamoxifen injection, followed by a 2-day rest period, and then an additional 5 days of tamoxifen injection, until the experimental endpoint.

### Mouse organotypic cultures

Organotypic cultures from adult mouse brains were prepared as previously described (Priego et al., 2018). In brief, brains were dissected in HBSS supplemented with HEPES (2.5 mM, pH 7.4), CaCl2 (1 mM), NaHCO3 (4 mM), MgCl2 (1mM) and D-glucose (30mM) embedded in 4% low-melting agarose (Lonza) preheated at 42°C. The embedded brains were cut into 250 mm slices using a vibratome (Leica) and each slice was divided in two pieces at the hemisphere. The slices were placed with flat spatulas on top of 0.8-μm pore membranes (Sigma-Aldrich) floating on slice culture media (DMEM, supplemented HBSS 27%, FBS 5%, D-Glucose (27 mM), L-glutamine (1 mM), and 100 IU/mL penicillin/streptomycin) and subjected to the corresponding procedure.

For compartment-controlled organotypic slice experiments, the same slicing and culture protocol was followed, but tissue was selectively resected to generate two conditions. In the tumour microenvironment-containing condition, slices were trimmed to retain both tumour tissue and an adjacent portion of the surrounding microenvironment. In the tumour-only condition, slices were trimmed to retain tumour tissue alone, excluding the adjacent tumour microenvironment. In these experiments, T-5224 was used at a final concentration of 0.5 μM both *in vitro* and *ex vivo*, with DMSO as a control.

For the pharmacological targeting experiments (T-5224 inhibitor), brains from wild-type C57BL/6 mice (6–8 weeks of age) bearing established brain metastases, generated by intracranial injection of BrM cells, were collected 10 days after inoculation. Brain slices were imaged at day 0 to confirm the presence of established metastases by bioluminescence imaging (BLI) in an IVIS instrument (PerkinElmer) after addition of D-luciferin (3 mg/mL; Syd Labs) to the culture medium. Slices were randomized according to the initial BLI signal, and slices with non-detectable signal were excluded. Selected slices were then maintained in culture and treated with T-5224 (HY-12270, MedChemExpress), which was prepared as a 100 mM stock solution in DMSO and used at a final concentration of 10 μM for 7 days. Control slices received the corresponding vehicle (DMSO). Brain slices were re-imaged after 7 days of treatment, and average radiance (photons/sec/cm²/steradian) from each slice was quantified using Living Image software, version 4.5. Growth rate was calculated as the fold change in BLI signal at day 7 relative to day 0 and normalized to the values obtained from control slices. At the end of the experiment, brain slices were fixed overnight in 4% paraformaldehyde (PFA) at 4 °C and processed for immunofluorescence.

### Middle Cerebral Artery Occlusion in mice

Distal permanent occlusion of the middle cerebral artery (MCA) was performed as previously described^44^. In brief, mice were anesthetized with 4% sevoflurane in a mixture of 80% air/20% oxygen, while body temperature was maintained at 37°C using a heating pad throughout the surgical procedure. Ophthalmic gel was placed on the eyes to prevent corneal drying. Buprenorphine (0.3 mg/kg, subcutaneously) was used for analgesia.

A scalp incision was made, and the left temporal muscle was dissected. The area between the zygomatic arch and the squamous bone was thinned using a high-speed drill and cooled with saline. The MCA was visualized with a stereomicroscope (PZMIV, World Precision Instruments, USA) and a thin bone layer over the MCA was carefully removed. A piece of Whatman filter paper soaked in freshly prepared 20% ferric chloride was placed over the intact dura mater covering the MCA for 10 minutes and then removed to allow thrombus formation.

### Infarct volume measurement

Magnetic resonance imaging (MRI) was performed 48 hours after MCAO using a T2 MRI scanner (7-Tesla MRI, Agilent-Varian, CNIC). Mice were sedated with 2% sevoflurane and body temperature was maintained with an airflow system. The whole brain was scanned in coronal sections and seven consecutive sections spanning from bregma 2.5 to bregma -2.5 were analysed using Fiji software. The areas of the contralateral hemisphere, ipsilateral hemisphere and infarct were measured. The oedema index was calculated using the following formula: Oedema= (ΣIpsilateral hemisphere area/ΣContralateral Hemisphere Area). Infarct volume was calculated using the following formula: Infact volume (%) = [Σ(Infarct Area/Oedema)*100]/ΣContralateral Hemisphere Area.

### Open Field test

Three days prior to testing, animals were handled daily by the experimenter in the behavioural testing room. Each animal was placed in the centre of the open field arena (40 cm x 40 cm x 70 cm) and its behaviour was recorded for 10 minutes using an overhead camera. The recordings were analysed using AnyMaze software. Total distance travelled, mean speed and time in the centre, inner zone and periphery were analysed.

### Cell Culture

Brain metastatic cell lines (BrM) from human and mouse origins have been previously described. Human H2030-BrM3 (abbreviated as H2030-BrM) cells were cultured in RPMI1640 media supplemented with 10% FBS, 100 IU/mL penicillin/streptomycin, 1 mg/mL amphotericin B and 2 mM L-glutamine. Murine B16/F10-BrM7^9^ (abbreviated as B16/F10-BrM) cells were cultured in DMEM media supplemented with 10% FBS, 100 IU/mL penicillin/streptomycin, 1 mg/mL amphotericin B and 2 mM L-glutamine. Murine 482N1 cells were cultured in DMEM medium supplemented with 10% FBS, 100IU.ml^−1^ penicillin/streptomycin, and 1 mg.ml^−1^ amphotericin B. Murine E0771-BrM cells were cultured in RPMI 1640 medium supplemented with 10 % FBS, 100 IU.ml^−1^ penicillin/streptomycin, 1 mg.ml^−1^ amphotericin B, and 1 % hepes. All cell lines were tested negative for Mycoplasma. All the cell lines harbored a plasmid expressing luciferase under the PGK promoter.

For the *in vitro* inhibition assay, 1.000 E0771-BrM cells per well were seeded in black 96 well-plates (Fisher Scientific, ref. 08-772-225). Plates were scanned by the Opera Phenix Plus High-Content Screening (HCS) System (Perking Elmer) 12 hours, 5 days and 7 days after seeding. Cells were automatically counted using Harmony v.5.2 software.

### Immunofluorescence and immunohistochemistry

Tissue for free-floating immunofluorescence was processed after overnight fixation at 4°C with 4% paraformaldehyde (PFA). Mice brains were processed, and mouse slices were performed by using a sliding microtome (80 mm, Thermo Fisher Scientific) or vibratome (250 mm). Slices were blocked using a blocking solution composed by 10% NGS, 2% BSA and a 0.25% Triton X-100 in PBS for 2h at RT. Primary antibodies were incubated at 4°C overnight in the blocking solution and the following day 30 min at RT. The appropriate fluorochrome-conjugated secondary antibody was added in blocking solution and incubated for 2 h at RT after extensive washing with PBS-Triton 0.25%. After extensive washing with PBS-Triton 0.25%, using DAPI (1 mg/mL, Sigma-Aldrich) for 7 min at RT nuclei were stained. After extensive washing with PBS, slices were mounted on glass slides using Fluoromount-G mounting medium (SouthernBiotech).

**Table 1:**
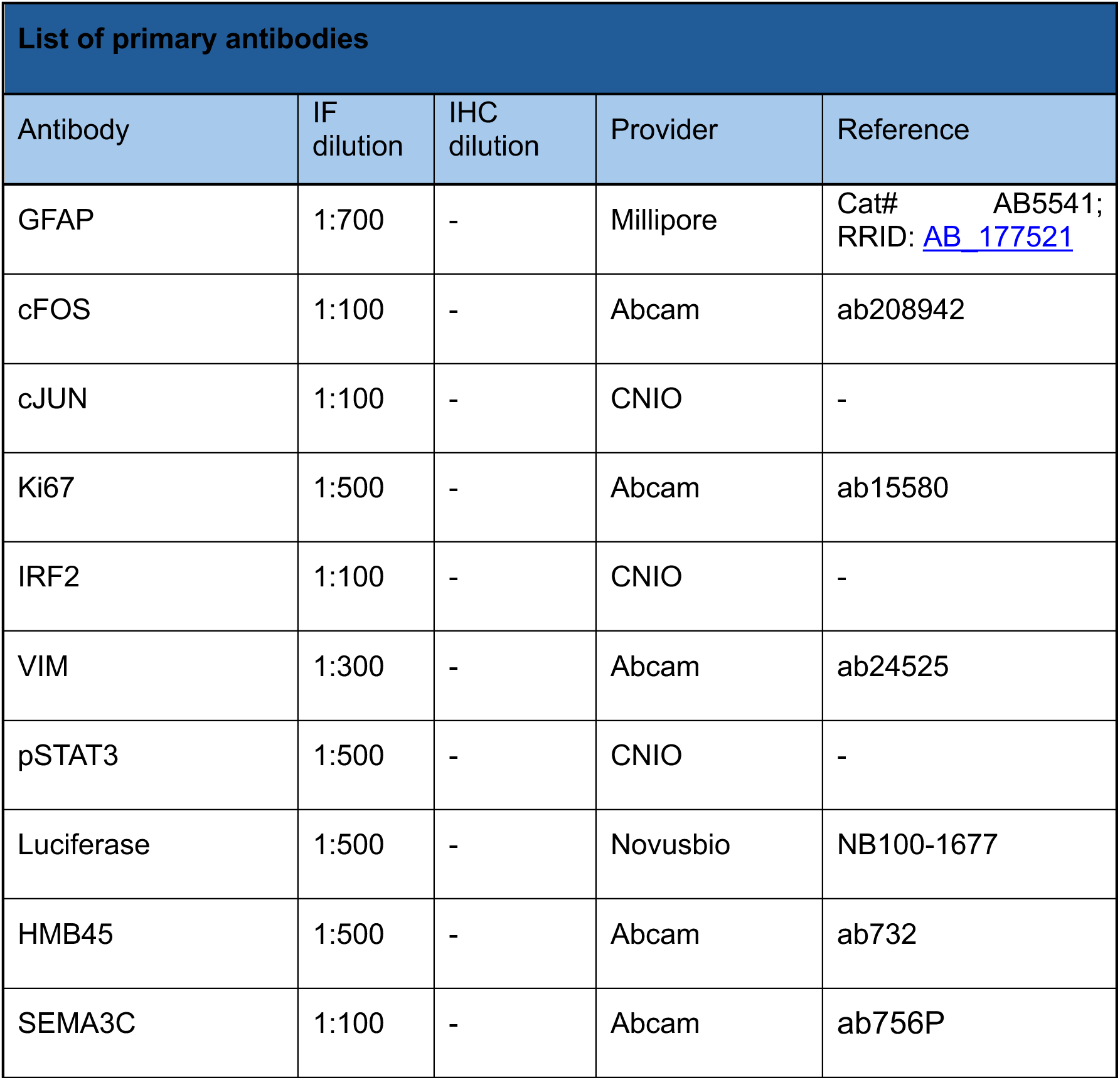
List of primary antibodies.

| List of primary antibodies |  |  |  |  |
|---|---|---|---|---|
| Antibody | IF dilution | IHC dilution | Provider | Reference |
| GFAP | 1:700 | - | Millipore | Cat# AB5541; RRID: AB_177521 |
| cFOS | 1:100 | - | Abcam | ab208942 |
| cJUN | 1:100 | - | CNIO | - |
| Ki67 | 1:500 | - | Abcam | ab15580 |
| IRF2 | 1:100 | - | CNIO | - |
| VIM | 1:300 | - | Abcam | ab24525 |
| pSTAT3 | 1:500 | - | CNIO | - |
| Luciferase | 1:500 | - | Novusbio | NB100-1677 |
| HMB45 | 1:500 | - | Abcam | ab732 |
| SEMA3C | 1:100 | - | Abcam | ab756P |

**Table 2:**
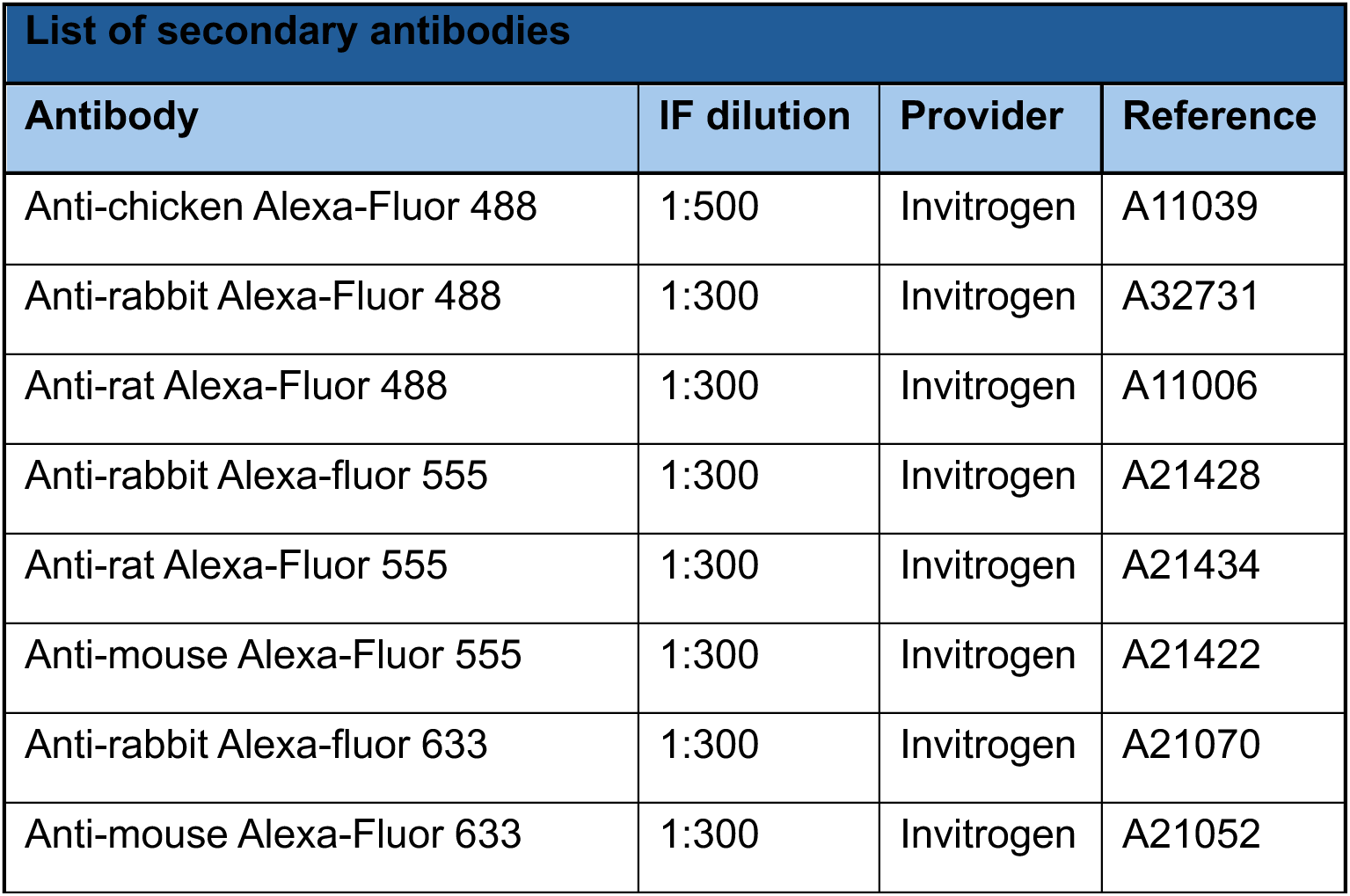
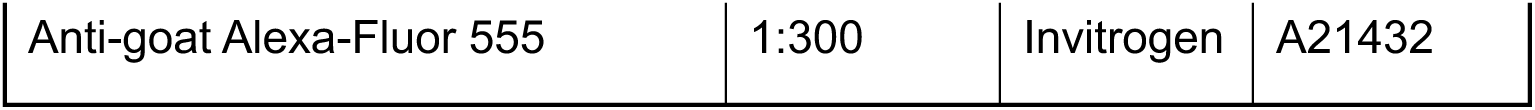
List of secondary antibodies.

| List of secondary antibodies |  |  |  |
|---|---|---|---|
| **Antibody** | **IF dilution** | **Provider** | **Reference** |
| Anti-chicken Alexa-Fluor 488 | 1:500 | Invitrogen | A11039 |
| Anti-rabbit Alexa-Fluor 488 | 1:300 | Invitrogen | A32731 |
| Anti-rat Alexa-Fluor 488 | 1:300 | Invitrogen | A11006 |
| Anti-rabbit Alexa-fluor 555 | 1:300 | Invitrogen | A21428 |
| Anti-rat Alexa-Fluor 555 | 1:300 | Invitrogen | A21434 |
| Anti-mouse Alexa-Fluor 555 | 1:300 | Invitrogen | A21422 |
| Anti-rabbit Alexa-fluor 633 | 1:300 | Invitrogen | A21070 |
| Anti-mouse Alexa-Fluor 633 | 1:300 | Invitrogen | A21052 |
| Anti-goat Alexa-Fluor 555 | 1:300 | Invitrogen | A21432 |

Immunohistochemistry of paraffin embedded tissues was performed at the CNIO Histopathology Core Facility. For the different staining methods, the slides were deparaffinized in xylene and rehydrated by a graded ethanol series to water. Several immunohistochemical reactions were performed on an automated immunostaining platform (Autostainer Link 48, Agilent; Discovery XT-ULTRA, Roche/Ventana). First, antigen retrieval was performed with the appropriate pH buffer and endogenous peroxidase was blocked (3% hydrogen peroxide). The slides were then incubated with the appropriate primary antibody for single or double staining. Following the primary antibody, the slides were incubated with appropriate secondary antibodies and with horseradish peroxidase (HRP)–conjugated visualization systems when needed.

The immunohistochemical reaction was revealed using ChromoMap DAB, DISCOVERY Purple, or Teal Kit (Roche/Ventana). Nuclei were counterstained with hematoxylin. Finally, slides were dehydrated, rinsed, and mounted for microscopic evaluation. Positive controls for primary antibodies were included in each staining series.

### Multiplex immunohistochemistry

Multiplex immunohistochemistry was carried out with OPAL technology (Quanterix Corp.) on the Ventana Discovery Ultra platform (Ventana Medical Systems), following a previously described protocol^45^. In brief, formalin-fixed, paraffin-embedded (FFPE) brain metastasis specimens were deparaffinized and underwent heat-induced antigen retrieval. Tissue sections were subsequently incubated with a primary antibody, followed by an HRP-conjugated secondary antibody. After addition of tyramide signal amplification (TSA) fluorophores (Quanterix Corp.), HRP converted the TSA molecules into reactive radicals, which covalently attached to tyrosine residues at the epitope. Antibody complexes were then removed by heat-mediated stripping, enabling the next staining round. This process was conducted five times in total, each round using different primary antibodies and TSA fluorophores with distinct emission spectra. The staining procedure was finalized with DAPI counterstaining (Sigma Aldrich), and slides were mounted using Fluoromount-G® medium (SouthernBiotech). Stained slides were stored protected from light at 4 °C until image acquisition.

After whole-slide scanning at 100x magnification using the PhenoImager HT 2.0 platform (Quanterix Corp.), regions of interest were defined in Phenochart™ software (Quanterix Corp.). Multispectral images (MSIs) were acquired at 200x magnification and spectrally unmixed with inForm software (Quanterix Corp.). The resulting MSIs were exported as multichannel TIFF files and stitched using QuPath software^46^. Representative images were generated with QuPath and Inkscape.

**Table 3:**
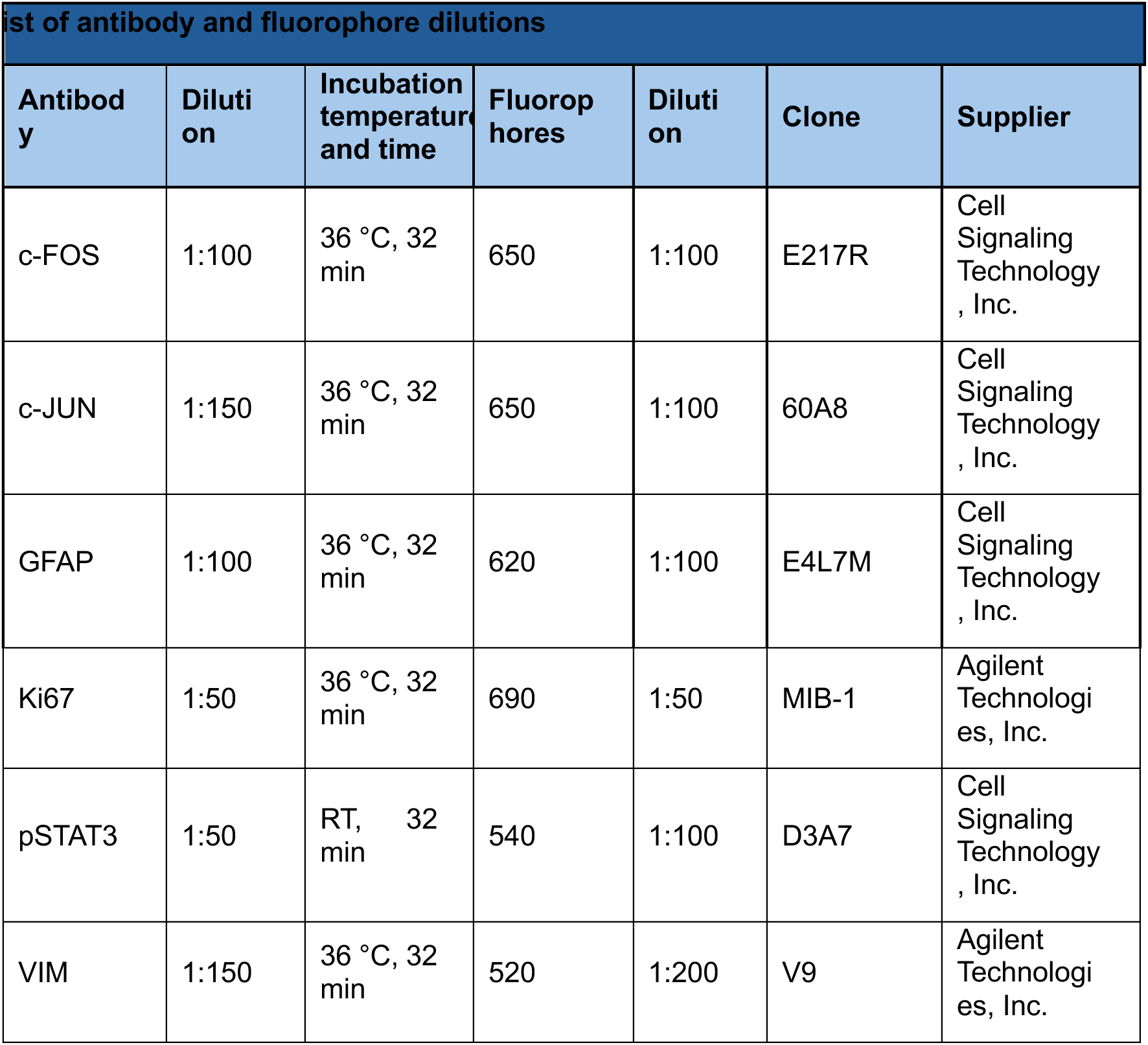
List of antibodies and fluorophore dilutions for multiplex immunohistochemistry.

### Image acquisition and analysis

Immunofluorescence images were acquired using a Leica Thunder imager (5x) and SP8, SP5 up-right confocal microscopes using 5x, 10x, 20x and 63x objectives and analyzed with Fiji software v2.7. Quantification and colocalization analyses were performed semiautomatically using groovy scripts ran on Fiji software v2.7. Immunohistochemistry images were captured with Zen Blue Software v3.1 (Zeiss). DAB immunohistochemistry intensity was measured using QuPath v.0.4.2.

### Processing of mouse brains for single-cell RNA sequencing

C57BL/6 mouse brains from were extracted in pre-cooled 1× D-PBS and processed using the Adult Brain Dissociation Kit (Miltenyi, cat. no. 130-107-677) with gentleMACS C Tubes (Miltenyi, cat. no. 130-093-237) and the gentleMACS Octo Dissociator (Miltenyi, cat. no. 130-096-427). The resulting cell suspension was filtered through a 70-μm strainer and centrifuged at 300g for 10 min at 4 °C. For myelin removal, the protocol described by Korin et al. was followed. Briefly, the cell pellet was resuspended in 7 mL of RPMI1640 at room temperature, and 3 mL of SIP solution (Stock Isotonic Percoll, Sigma-Aldrich, cat. no. GE17-0891-02) was added to each tube with gentle mixing. The resulting 30% (vol/vol) Percoll/cell mixture was carefully layered on top of 2 mL of 70% (vol/vol) SIP in 1× PBS. Samples were centrifuged at 500g for 30 min at 18 °C with minimal deceleration. The top myelin layer was removed using a 10-mL pipette, and the remaining cell suspension was centrifuged again at 500g for 7 min at 18 °C. The supernatant was discarded, and cells were resuspended in cold D-PBS containing 0.5% BSA for staining. Cell suspensions were magnetically labeled with anti-ACSA2 microbeads (Miltenyi, cat. no. 130-097-678), and enrichment of glial populations was verified by flow cytometry (BD FACSCanto II) using anti-ACSA2-PE (1:100, Miltenyi, cat. no. 130-123-284)^28^. For dead cell removal and washing before single-cell sequencing, Debris Removal Solution (Miltenyi, cat. no. 130-109-398) was used. The live-cell fraction was then centrifuged at 300g for 10 min, washed, and finally resuspended in 1× PBS containing 0.04% BSA at a concentration of 7 × 10^5 cells/mL, and kept on ice. Cells suspended in PBS/BSA were assessed for viability and checked to ensure the absence of debris and aggregates. The cell suspension was then loaded onto a 10x Chromium Single Cell Controller Chip B (10x Genomics).

For transgenic model GFAP-CRE^ERT2;^ Ai14(tdTomato) only peritumoral area was processed and DSP-fixation was applied before tissue dissociation following the protocol described by Jimenez-Garcia et al. After fixation, samples were processed according to the workflow described above, including tissue dissociation, myelin removal and washing steps. Unlike C57BL/6 samples, which were subjected to ACSA2-based enrichment, transgenic samples were purified by fluorescence-activated cell sorting of tdTomato-positive cells. Final cell suspensions were resuspended in 1× PBS containing 0.04% BSA, assessed for viability and absence of debris and aggregates, and loaded onto a 10x Chromium Single Cell Controller Chip B (10x Genomics) according to the manufacturer’s instructions (Chromium Single Cell 3′ GEM, Library & Gel Bead Kit v3, PN-1000075).

| List of flow cytometry antibodies |  |  |  |
| Antibody | IF dilution | Provider | Reference |
|---|---|---|---|
| ACSA2-PE | 1:50 | Milteny |  |

### Single cell RNA sequencing library preparation and preprocessing

ACSA2⁺ fraction cells were processed for single-cell RNA sequencing library preparation using the Chromium Next GEM Single Cell 3′ GEM, Library & Gel Bead Kit v3.1 (10x Genomics, PN-1000121), following the manufacturer’s instructions. The protocol included GEM generation, barcoding, GEM-RT clean-up, cDNA amplification, and gene expression library construction. Libraries were sequenced on an Illumina NextSeq 550 instrument equipped with v2.5 reagent kits, using a paired-end configuration with 28 bp for Read 1 and 56 bp for Read 2. Primary data processing, including image analysis, per-cycle base calling, and quality score assignment, was performed using Real Time Analysis software (RTA v2, Illumina). BCL files were converted to FASTQ format using bcl2fastq2 within the Local Run Manager RNA Fusion Analysis Module v2 (Illumina).

FACS-purified tdTomato⁺ cells were processed using the Chromium Next GEM Single Cell 5′ Kit v2 (10x Genomics), following the manufacturer’s instructions. Briefly, each cell suspension was loaded into one port of a Chromium Next GEM Chip K (10x Genomics). Single cells were encapsulated into emulsion droplets using the Chromium Controller (10x Genomics). ScRNA-seq libraries were prepared using the Chromium Next GEM Single Cell 5’ Kit v2 (10x Genomics) following the manufacturer instructions and each library was amplified using a SureCycler 8800 thermal cycler (Agilent Technologies). The average size of each library was then calculated using a High sensitivity DNA chip on a 2100 Bioanalyzer (Agilent Technologies) and the concentration was determined using the Qubit fluorometer (Thermofisher). Individual libraries were diluted to 10 nM and pooled for sequencing. Library pool was loaded at 700 pM onto a P3 flow cell (100 cycles) of the NextSeq 2000 (Illumina) in paired-end configuration (28bp Read1, 10bp Index1, 10pb Index 2 and 90bp Read2). FastQ files for each sample were obtained using cellranger mkfastq pipeline (10x Genomics).

Raw sequencing reads were aligned to the mouse reference genome (GRCm38, GENCODE vM25 annotation) using STARsolo (v2.7.8a). Downstream analyses, including quality control, normalization, dimensionality reduction, clustering and marker gene identification, were performed using Seurat (v5.1.0). Astrocyte identities were assigned based on previously published signatures from Zeisel, Habib, Liddelow, Batiuk and Martirosyan and colleagues, following the annotation strategy described by Priego et al. Following quality-control filtering, 8,670 astrocytes were retained for analysis, including 592 cells from 482N1-early-model technical replicate 1, 884 from 482N1early model technical replicate 2, 306 from E0771-BrM late model technical replicate 1, 564 from E0771-BrM late model technical replicate 2, 1,158 from 482N1 late model, 2,158 from B16/F10-BrM and 3,008 from healthy sample. Cells contained a mean of 1,867 detected genes and 4,122 UMIs per cell (medians: 1,747 genes and 3,187 UMIs per cell, respectively).

### Single cell RNA sequencing downstream processing, integration and annotation

For the early-stage dataset, the two technical replicates were initially analyzed independently. Cells expressing fewer than 200 genes or more than 3,000 genes, as well as cells with more than 25% of unique molecular identifiers (UMIs) derived from mitochondrial genes, were excluded. After quality filtering, 19,331 genes and 876 cells were retained from technical replicate 1, while 19,872 genes and 1,172 cells were retained from technical replicate 2. Data normalization was performed using the NormalizeData function in Seurat with default parameters. Highly variable genes were identified with FindVariableFeatures, selecting the top 2,000 variable features. Dimensionality reduction was performed by principal component analysis (PCA), followed by uniform manifold approximation and projection (UMAP) using the top 50 principal components. Cell clustering was performed using FindClusters.

Major cell populations within the tumour microenvironment were annotated separately in each sample using canonical marker genes. In the early-stage compartment, both technical replicates consistently identified the same major cell classes, including endothelial cells (*Pecam1, Cldn5*), neurons (*F5*), microglia/macrophages (*Cd68, C1qa, C1qb, C1qc, Tmem119*) and astrocytes (*Gfap, Aqp4, S100b*). To further validate these annotations, the early-stage dataset was integrated with previously annotated late-stage and healthy datasets representing the tumour microenvironment.

The two early technical replicates were subsequently integrated using Seurat FindIntegrationAnchors and IntegrateData. Given the high concordance between replicates, the integrated dataset was used for downstream analyses. Astrocytes were then subset using Seurat for dedicated analysis. Within this astrocyte-only dataset, normalization, variable feature selection, PCA, clustering and UMAP were repeated. Multiple clustering resolutions were tested, and the final resolution (RNA_snn_res.0.2) was selected based on biological interpretability and separation of functionally distinct astrocyte populations.

Astrocyte identities were assigned using a signature-enrichment strategy based on curated marker genes and published reference signatures from Zeisel, Habib, Liddelow, Batiuk and Martirosyan and colleagues. Signature scores were calculated using Seurat AddModuleScore, allowing each cluster to be evaluated according to enrichment for previously described astrocyte programs. To minimize contamination by non-astrocyte populations, the early astrocyte dataset was cross-referenced with the late-stage tumour microenvironment dataset through anchor-based integration. Ambiguous cells were examined using canonical marker gene expression and cluster context, and cells clustering with endothelial, neuronal or microglia/macrophage populations were excluded. After removal of contaminating populations, the astrocyte dataset was re-normalized and re-clustered, yielding a final early astrocyte dataset composed of 11 clusters.

For cross-condition comparisons, astrocyte datasets from healthy, early and late conditions were integrated and clustering resolution was iteratively optimized, resulting in a final integrated resolution of integrated_snn_res.0.2. Cluster-specific marker genes were identified using FindAllMarkers and FindMarkers. To compare transcriptional similarity across cluster-condition combinations, the top 25 marker genes for each cluster-condition pair were extracted and used to generate enrichment profiles. Weighted mean enrichment scores were calculated using run_wmean, allowing systematic comparison of shared transcriptional programs across conditions.

Gene set enrichment analysis (GSEA) was performed using GSEA Desktop v4.3.3 in preranked mode with the MSigDB mouse collections mh.all.v2024.1.Mm.symbols.gmt, m2.all.v2024.1.Mm.symbols.gmt and m5.all.v2024.1.Mm.symbols.gmt. The chip annotation file Mouse_Ensembl_Gene_ID_MSigDB.v2024.1.Mm.chip was used. Analyses were run with the weighted enrichment statistic, without collapsing to gene symbols, and with gene-set size limits of 15 to 500 genes. Ranked gene lists were generated from Seurat differential expression outputs, ordered by average log2 fold change and filtered at adjusted *P* < 0.05. GSEA was performed for early versus control comparisons to characterize early astrocyte programs and for early versus late comparisons to identify stage-dependent transcriptional changes. Pathways with false discovery rate (FDR) q values ≤ 0.05 were considered significant.

### Analysis of our AST states across publicly accessible human scRNA-seq datasets

To assess the conservation of murine astrocyte states in human disease, we re-analyzed a publicly available single-cell RNA-seq dataset of human NSCLC brain metastases^26^. Following the annotation strategy described above for mouse datasets, major cell populations were first manually annotated using canonical marker genes, and the astrocyte compartment was subsequently subset for dedicated downstream analysis.

Within the human astrocyte subset, data were normalized, dimensionality reduction was performed, and clustering resolution was selected in an unbiased manner based on cluster separation and biological interpretability. UMAP was used for visualization of the human astrocyte compartment.

To compare human astrocyte populations with the murine AST states identified in our study, we applied the same signature-enrichment strategy used for mouse astrocyte annotation. Murine AST state signatures were projected onto the human astrocyte subset using enrichment scoring, and human clusters were evaluated according to their relative enrichment for each murine programme. Human astrocyte clusters showing shared marker-gene structure and concordant enrichment with murine AST states were annotated as corresponding human astrocyte states (hASTs).

Cross-species correspondence between human and mouse astrocyte states was visualized as a dot plot summarizing the relative enrichment/overlap between each human–mouse state pair. Dot colour represents the relative enrichment or correlation score, and dot size reflects the proportion of shared marker representation. This analysis enabled identification of conserved murine-defined astrocyte programs in human NSCLC brain metastases while also revealing additional heterogeneity within the human astrocyte compartment.

### Visium library preparation and sequencing

Spatial transcriptomic profiling was performed using the Visium Spatial Gene Expression platform (10x Genomics) according to the manufacturer’s instructions. OCT-embedded mouse brain samples were sectioned at 10 μm using a cryostat (Leica CM1950). Tissue sections were mounted onto Visium slides, H&E stained, and imaged using a NanoZoomer S60 scanner (Hamamatsu) to assess tissue morphology and quality. Permeabilization time was optimized using the Visium Tissue Optimization workflow and set to 6 min. Reverse transcription, second-strand synthesis, cDNA denaturation, amplification and library construction were performed according to the Visium Spatial Gene Expression User Guide (10x Genomics). Spatially barcoded full-length cDNAs were amplified by PCR for 16 or 18 cycles depending on initial cDNA yield, as determined by qPCR. Indexed libraries were generated by end repair, A-tailing, adaptor ligation and sample index PCR, and library quality was assessed using an Agilent 2100 BioAnalyzer. Libraries were sequenced on an Illumina NovaSeq 6000 at a depth of approximately 200,000 reads per spot.

### Visium data preprocessing, clustering and annotation

Sequencing data were processed with Space Ranger (v1.3.0) using the mouse reference genome to generate filtered feature-barcode matrices. These matrices were used for downstream analyses in Seurat (v4.1.1). Spots with fewer than 500 UMIs or lacking tissue coverage were excluded. Data were log-normalized using NormalizeData with a scale factor of 10.000. Highly variable genes were identified with FindVariableFeatures, selecting the top 3.000 features. Data were then scaled using ScaleData, and principal component analysis (PCA) was performed. The top 30 principal components were used for downstream analyses.

For clustering, a nearest-neighbor graph was constructed with FindNeighbors, and clusters were identified using the Leiden algorithm implemented in FindClusters. A slide-specific resolution ranging from 0.4 to 0.6 was selected according to tissue structure and cluster separation. Differentially expressed genes for each cluster were identified using FindAllMarkers with the Wilcoxon rank-sum test. Clusters were manually annotated to anatomical brain regions using curated marker genes and the Allen Brain Atlas as reference.

### Spatial deconvolution and downstream analyses

To estimate cell-type composition in each Visium spot, we used a single-cell reference from mouse brains with or without B16/F10-BrM intracranial metastases (Priego et al., 2025). Deconvolution was performed using the seeded non-negative matrix factorization regression framework implemented in SPOTlight (v1.0.2).

Pathway and transcription factor activities were inferred using decoupleR (v2.2.2). Fourteen mouse pathways from PROGENy and the DoRothEA gene regulatory network (confidence levels A–C) were used as prior knowledge resources. Activity scores were estimated using the weighted mean method (run_wmean) on normalized spot-level expression values.

To identify gene programs shared across time points and conditions, topic modeling was applied to the raw count matrix using fit_topic_model from the fastTopics package (v1.0.0). Topic models were fitted jointly across all samples using 20 topics (K = 20). Topic-associated genes were defined using the elbow point identified with the uik function from the inflection package (v1.3.5). These topic-derived gene sets were subsequently analyzed by gene set enrichment analysis using fgsea (v1.20.0) and mouse gene sets retrieved through the msigdbr package (v7.5.1). Mitochondrial and ribosomal genes were excluded from these analyses. Topics were further annotated using de_analysis, which implements grade-of-membership differential expression analysis by weighting each spot according to its topic contribution.

### Human sample collection

Human brain metastasis were collected by CNIO Biobank from different hospitals of our consortium: Hospital Universitario 12 de Octubre, Hospital Universitario La Princesa, Hospital Universitario de Bellvitge, Hospital Universitario de Burgos, Hospital Universitario de Navarra, Complejo Hospitalario Universitario de Vigo. All the studies were conducted in accordance with recognized ethical guidelines (Declaration of Helsinki) and were approved by the Research Ethics Committee of the Carlos III Health Institute (ref. CEI PI 91_2023; ref. CEI-PI 63_2023; ref. CEI PI 25_2020-v2_Amendment 2021, ref. CEI PI 91_2023, ref. CEI PI 25_2020, CEI PI 51_2023 and CEI PI 63_2023). Written informed consent from each donor was collected. Comprehensive clinical information is also collected by CNIO Biobank associated with the samples. The samples were transported to Spanish National Cancer Research Centre within 24 hours after surgical resection under controlled temperature conditions and standardized preanalytical procedures to ensure cohort homogeneity and sample quality.

### Purity-Stratified Survival Analysis

To evaluate the prognostic impact of specific transcriptomic signatures (FOS and FOS astrocyte gene sets) within distinct tumor microenvironments (TME), normalized and batch-corrected bulk RNA-seq count matrices from the brain metastasis RENACER cohort were preprocessed by applying a log2(x+1) transformation prior to calculating sample-wise enrichment scores via Gene Set Variation Analysis (GSVA; Gaussian cumulative distribution function, minimum gene set size = 3). For clinical modeling, overall survival (time from brain metastasis diagnosis to death or last follow-up) served as the primary endpoint, and covariates were systematically formatted: age at diagnosis was categorized into three brackets (< 50, 50–65, and > 65 years), clinical stage was collapsed into early (I–III) versus late (IV) disease, and rare primary tumor anatomies (n < 10) were pooled into an “Other” category with breast cancer designated as the statistical baseline. To effectively decouple intrinsic tumor biology from stromal background signals, the global cohort was evaluated both in its entirety (“All Data”) and stratified by the median tumor purity score to establish distinct “Tumor-rich/TME-low” (high purity samples) and “Stroma-rich/TME-high” (low purity samples) sub-cohorts. Within each specific sample stratification, patients were stratified into high- and low-expression for each transcriptomic signature separately tiers using symmetric, independently calculated percentile thresholds (median split, as well as top versus bottom 40%, 30%, 20%, and 10% extremes). Survival trajectories for these subsets were estimated using the Kaplan-Meier method and compared via log-rank tests, while independent prognostic value was determined by fitting multivariable Cox proportional hazards regression models adjusted for primary site, age, sex, and stage. Adjusted hazard ratios (HR) and 95% confidence intervals (CI) were ultimately extracted and visualized using custom dumbbell-style forest plots to systematically illustrate the spatial shift of signature-associated mortality risk across varying tumor purities.

### Quantification and statistics

Data were analyzed using GraphPad Prism 8 software (GraphPad Software). For comparisons between two experimental groups in datasets that followed a normal distribution, an unpaired or paired (depending on the data), two-tailed Student t test was used. For comparisons that did not follow a normal distribution, a Mann-Whitney U test were performed. For multiple comparisons, ANOVA test was performed. For survival curves, P values were obtained with log-rank (Mantel–Cox) two-sided tests. χ2 test was performed for the comparison of group proportions.

## Data availability and code

All data will be made available upon request to

scRNA-seq and spatial transcriptomics data have been deposited in GEO under accession number GSE337748, GSE337749, GSE337751, GSE337764.

Code used for data processing and downstream analyses is available at GitHub (https://github.com/ValienteLab/AST-BrM-atlas).

## Acknowledgements

We thank all members of the Brain Metastasis Group for critical discussion of the manuscript; the CNIO Core Facilities for their excellent assistance. Special thanks to patients and their families who donated valuable materials to RENACER and all site personnel, investigators, funders, and industry partners who supported the generation of the data within this study. This study was funded by AECC (LABAE19002VALI, PRYCO234528VALI; M.V.), Fundación Ramón Areces (CIVP19S8163; M.V. and CIVP20S10662; E.O-P and MJ.A.), ERC CoG (864759; M.V.), La Caixa Foundation (HR23-00051; M.V.), Instituto de Salud Carlos III (PT23/00065; E.O-P and MJ.A.), Federal Ministry of Research, Technology and Space and co-funded by the European Commission (01KT2304B; M.S.), Federal Ministry of Research, Technology and Space (03ZU2111AB; M.S), Spanish Ministry of Science, Innovation and Universities (PID2022-140616OB-I00; MA.M).

L.C-B. and L.C. were funded by CNIO Friends Postdoctoral Fellowships, L.A-E received funds from MINECO-Severo Ochoa PhD Fellowship (BES-2017-081995). MG-B. is supported by the Spanish Ministry of Science, Innovation and Universities through the State Program to Develop, Attract and Retain Talent (FPU2022-01155). C.P-P received support from Spanish Ministry of Science, Innovation and Universities (PRE2021-099443, FPI).

The CNIC is supported by the Instituto de Salud Carlos III (ISCIII), the MICIU and the Pro CNIC Foundation, and is a Severo Ochoa Center of Excellence (grant CEX2025-001658-S funded by MICIU AEI/10.13039/501100011033). This work acknowledges the use of ICTS-ReDIB, supported by the MICIU at BioImaC.

## Declaration of competing interests

M.V. declares research agreements with AstraZeneca. The rest of co-authors declares no conflict of interest.

## Notes

### Competing Interest Statement

M.V. declares research agreements with AstraZeneca and Trogenix. The rest of co-authors declares no conflict of interest.

