## Supplementary Figures (1-4) and Supplementary Tables (1-4) (legends) for "Recurrent astrocyte transcriptional states with spatial and functional specialization are induced by brain metastases independently of their primary tumor origin"

**Supplementary Figure 1:** Aditional characterization of recurrent astrocyte states across primary tumor origins by single cell RNA-seq and spatial transcriptomics.

**Supplementary Figure 2:** Early brain metastasis profiling identifies transcriptionally distinct astrocyte states.

**Supplementary Figure 3:** Hopx deficiency improves recovery after ischemic injury and summarizes context-dependent roles of AST-BrM-associated programs.

**Supplementary Figure 4:** T-5224 has minimal in vitro effects but impairs brain metastatic growth ex vivo.

**Supplementary Tables (1-4):**

**Supplementary Table 1.** Differentially expressed genes between AST-BrM and homeostatic-like astrocyte states.

**Supplementary Table 2:** Top 50 differentially expressed genes for each AST-BrM cluster.

**Supplementary Table 3:** Filtered differentially expressed genes for each early AST-BrM cluster.

**Supplementary Table 4:** RENACER IDs used for Multiplex immunofluorescence validation in human BrM.

### Supplementary Figures

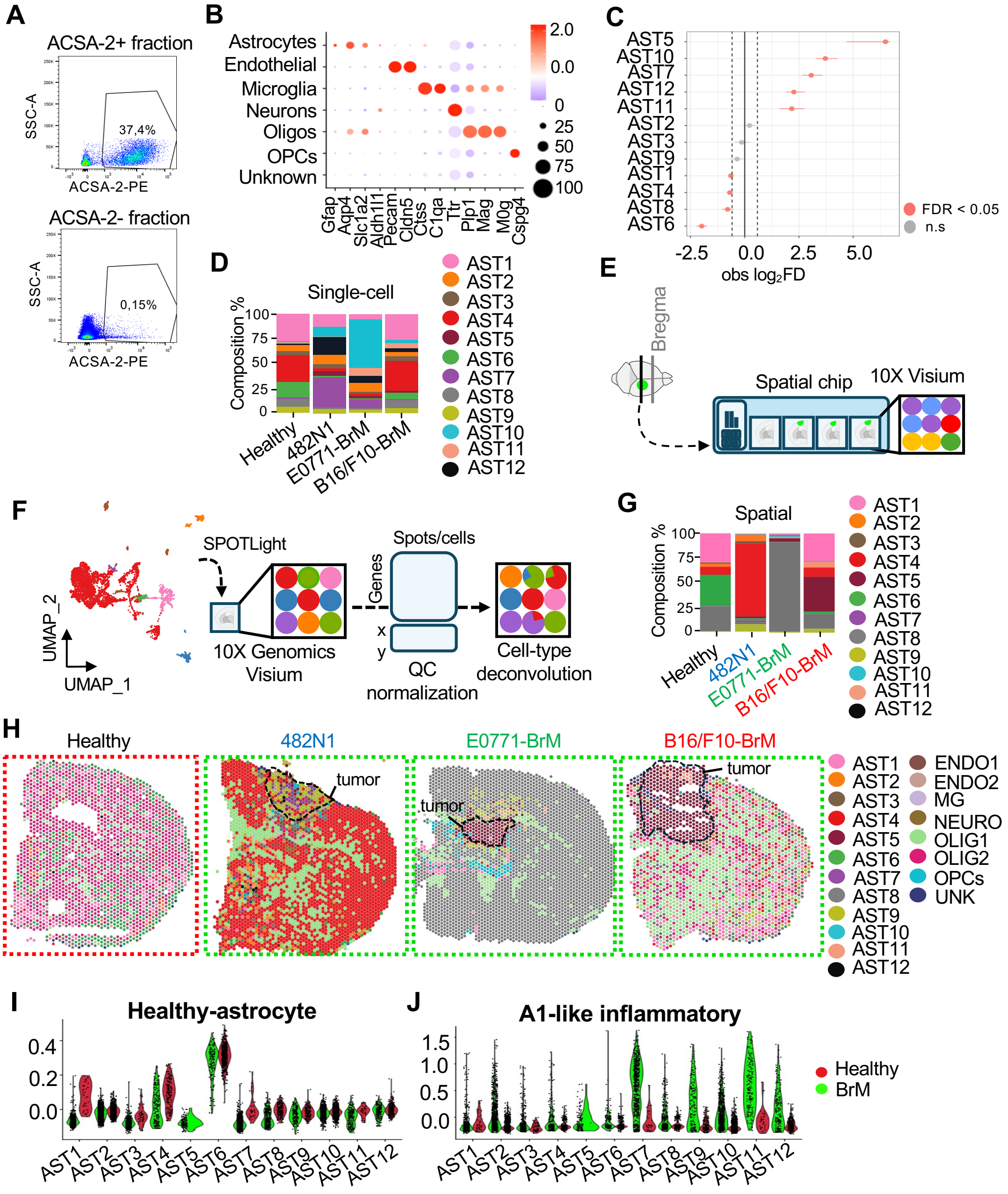


**Supplementary Figure 1: Aditional characterization of recurrent astrocyte states across primary tumor origins by single cell RNA-seq and spatial transcriptomics. A.** Representative flow cytometry plots showing the ACSA2-positive and ACSA2-negative fractions used for enrichment prior to single-cell RNA sequencing. **B.** Dot plot of canonical marker genes of the major cell classes captured in the dataset. Dot size indicates the proportion of cells expressing each marker, and color indicates scaled average expression: blue, low expression and red, high expression. **C.** Differential abundance analysis of astrocyte states between healthy and BrM conditions, shown as observed log2 fold change (obs log2FD). Red dots indicate significantly enriched states (FDR < 0.05), whereas grey dots indicate non-significant changes. **D.** Barplot showing the relative proportions of astrocyte states across healthy tissue and each BrM model in the single-cell dataset. **E.** Schematic overview of the spatial transcriptomic workflow. Coronal brain sections were collected at the indicated bregma level (0- 2), placed on 10x Visium spatial gene expression slides, and processed for downstream spatial transcriptomic analysis. **F.** Schematic representation of the deconvolution strategy. Cell-state signatures derived from the single-cell RNA-seq reference were projected onto 10x Visium spots with SPOTlight following quality control and normalization to infer spot-level cell-type composition. **G.** Relative proportions of astrocyte states (AST1–AST12) across healthy tissue and the different tumor-origin models, as inferred from spatial transcriptomic deconvolution. Although AST4 appears enriched in the lung model and AST8 in the breast model, spatial mapping (Suppl. Fig. 2E) shows that both states are broadly distributed in these tissues rather than selectively localized to the tumour interface. **H.** Spatial deconvolution maps showing the dominant (highest-proportion) cell-state assignment per spot across healthy brain and brain metastasis models. Tumor regions are outlined in black. Red dashed boxes indicate healthy samples, whereas green dashed boxes indicate brain metastasis models. **I.** Violin plots showing enrichment of the healthy-astrocyte signature from Zeisel et al., 2015 across the 12 astrocyte states in healthy and BrM conditions. Color legend in **J.** Violin plots showing enrichment of the A1-like inflammatory signature from Liddelow *et al.,* 2018 across the 12 astrocyte states in healthy and BrM conditions. Red color represents healthy condition whereas green color represents BrM models.

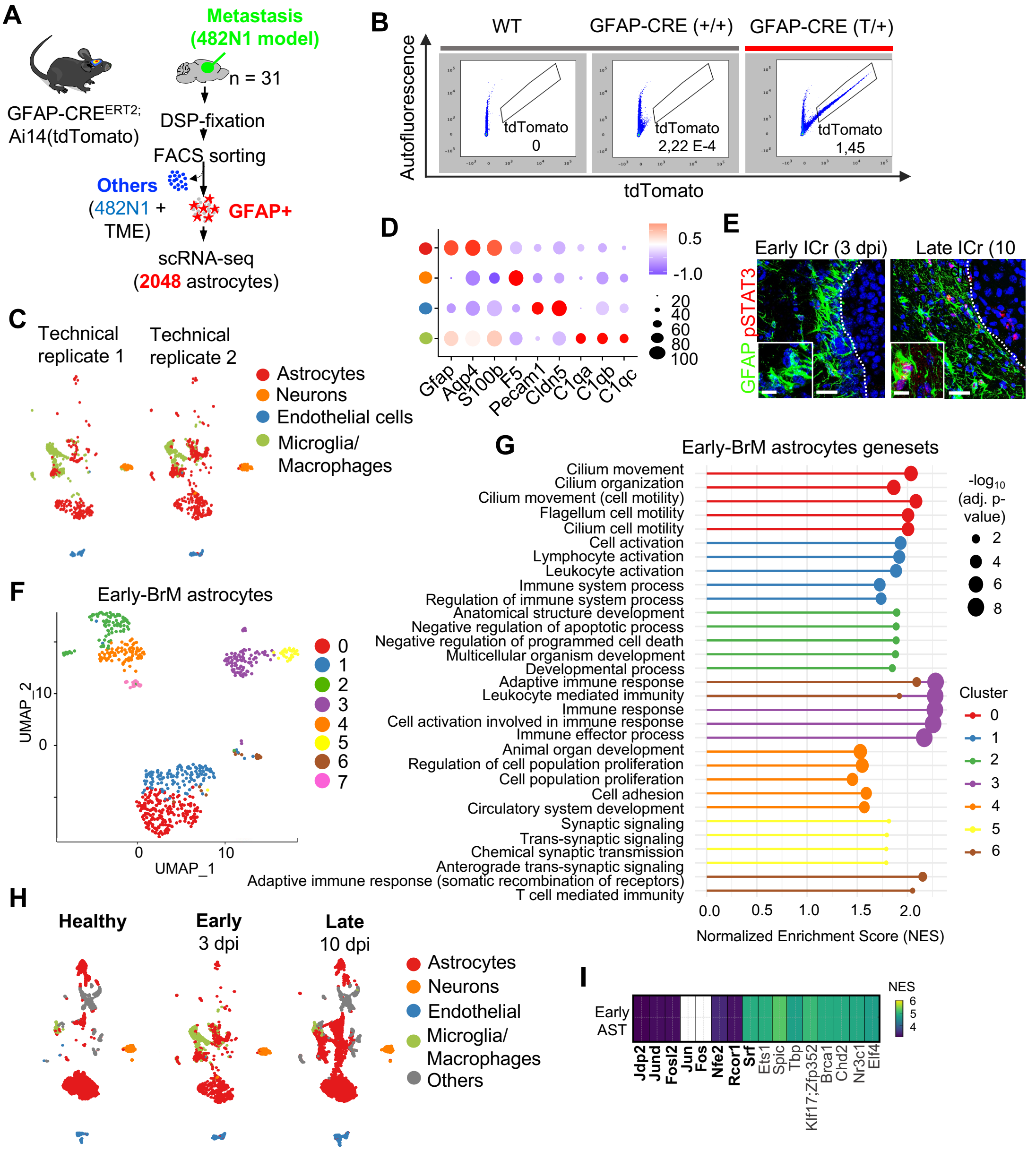


**Supplementary Figure 2: Early brain metastasis profiling identifies transcriptionally distinct astrocyte states**. **A.** Schematic overview of the experimental strategy. GFAP-CreERT2; Ai14(tdTomato) mice were injected intracranially with 482N1 cells, and brains were collected at early time points after metastasis induction (3 dpi). Following DSP fixation and FACS sorting, tdTomato$^{+}$ GFAP-lineage cells were isolated for scRNA-seq. A total of 2,048 astrocytes were retained for downstream analyses. **B.** Representative FACS gating strategy showing tdTomato fluorescence in WT and GFAP-Cre controls, and tdTomato$^{+}$ population in GFAP-CreERT2;Ai14(tdTomato) mice used for astrocyte enrichment. **C.** UMAP embeddings of two technical replicates from the early time point that were manually annotated independently. Cells colored by major cell type captured after sorting, including astrocytes, neurons, endothelial cells and microglia/macrophages. **D** Dot plot of canonical marker genes supporting cell-type annotation (astrocytes, neurons, endothelial cells, microglia/macrophages). Dot size indicates the fraction of cells expressing each gene; color indicates scaled average expression. Red for higher expression; blue for lower expression.

**E.** Immunofluorescence validation of reactive astrocyte activation across disease stages in the intracranial B16/F10-BrM model. Representative images of GFAP (green) and pSTAT3 (red) at early (3 dpi) and late (10 dpi) stages, showing increased $pSTAT3^+$ reactive astrocytes at late stage. **F.** Unbiased clustering of early BrM-associated astrocytes identifies transcriptionally distinct early astrocyte clusters (0–7) displayed on a UMAP embedding. **G.** Gene set enrichment analysis (GSEA) across early astrocyte clusters highlighting cluster-specific functional programs, including cilia-related gene sets, immune activation modules and synaptic-associated terms. Color denotes cluster identity; dot size reflects $-\log_{10}$ adjusted p-value; x-axis shows normalized enrichment score (NES). Cluster 7 has no significant hit potentially due to its low number of cells. **H.** UMAP across healthy, early and late timepoints for the major cell types identified. Colored by cell type. **I.** Summary of overlap across astrocyte gene sets/programs detected across early astrocytes after motif enrichment analysis.

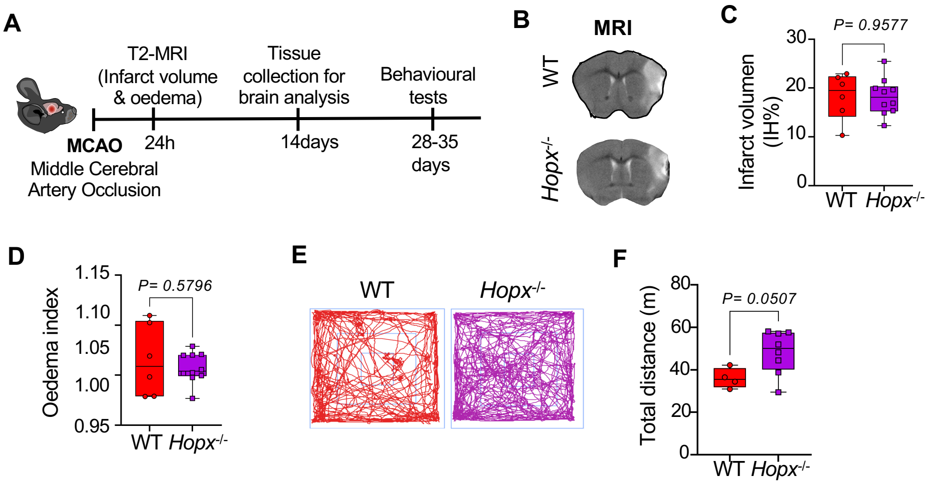


**Supplementary Figure 3: Hopx deficiency improves recovery after ischemic injury and summarizes context-dependent roles of AST-BrM-associated programs. A.** Experimental timeline of the middle cerebral artery occlusion (MCAO) model used to assess the role of Hopx in ischemic injury and recovery. T2-weighted MRI was performed 48 h after MCAO to quantify infarct volume and oedema, followed by tissue collection at 14 days and behavioural assessment at days 28–35. **B.** Representative Acute MRI assessment performed 48h after MCAO. **C.** Quantification of infarct volume in WT and Hopx $^{-/-}$ mice 24 h after MCAO. **D.** Quantification of oedema index in WT and Hopx $^{-/-}$ mice 24 h after MCAO. **E**. Trackplots from Open Field. **F.** Total distance travelled during behavioural testing in male WT and Hopx$^{-/-}$ mice.

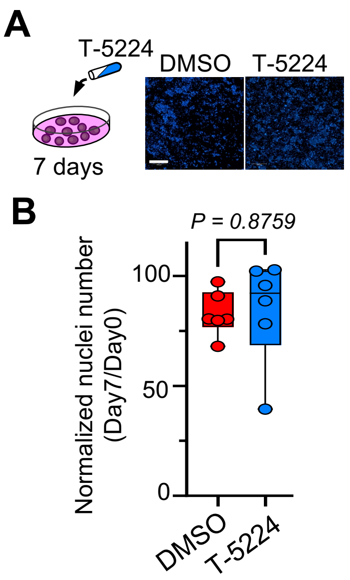


**Supplementary Figure 4: T-5224 has minimal in vitro effects but impairs brain metastatic growth ex vivo. A.** Experimental scheme of in vitro treatment with the AP-1 inhibitor T-5225 and representative images of BrM cells treated with DMSO or T-5224 for 7 days. Scale bar, 50 µm. **B.** Quantification of tumor cell number after DMSO or T-5224 treatment *in vitro*, showing no significant difference between conditions.
